# Structural basis of RNA-guided DNA integration by type I CRISPR-associated transposases

**DOI:** 10.64898/2026.05.18.725949

**Authors:** Giada Finocchio, Seraina Oberli, George Lampe, Michael Schmitz, Samuel H. Sternberg, Martin Jinek

## Abstract

CRISPR-associated transposases (CASTs) achieve site-specific DNA integration by coupling the RNA-guided targeting action of a nuclease-deficient CRISPR-Cas system with the assembly of a Tn7-like transpososome complex^1,2^. Understanding the detailed mechanisms of this elaborate process is paramount to engineering CAST systems into programmable genetic tools^3–6^. The type I-F *Pseudoalteromonas* CAST (*Pse*CAST) displays the highest activity in mammalian cells to date^7^ and has been the subject of extensive directed evolution^8^, but efforts to rationally engineer further improvements have been hampered by critical gaps in our understanding of transpososome assembly and activation^9^. Here we use cryo-EM structural analysis, validated by DNA transposition assays, to visualize the *Pse*CAST system in a series of functional states that define the stepwise mechanism of RNA-guided DNA integration. The structure of a target DNA-bound Cascade-TniQ-TnsC complex reveals that conformational changes induced by R-loop formation are coupled to target DNA stabilization and TnsC heptamerization, which in turn recruits the TnsAB transposase via conserved interactions with its C-terminal tail. Finally, the structure of the 1.2 MDa *Pse*CAST transpososome holocomplex reveals specific TnsC-TnsB and TnsB-target DNA interactions that drive allosteric remodelling of the TnsB catalytic site to activate donor DNA integration. Together, these findings establish a unified structural and mechanistic blueprint for RNA-guided DNA integration and lay the foundation for engineering next-generation DNA insertion systems for genome editing applications.

## INTRODUCTION

Transposons are pervasive, selfish DNA segments capable of mobilizing within and between genomes, which act as potent drivers of genome evolution and horizontal gene transfer^10–12^. CRISPR-associated transposons have evolved repeatedly via exaptation of minimal, nuclease-deficient CRISPR-Cas systems to achieve RNA-guided, site-specific DNA integration^13–15^, with emerging applications in genome editing. These systems operate through the concerted action of a CRISPR-based targeting module and a Tn7-like transposition machinery that comprises the TnsC AAA+ ATPase, the TnsB transposase, and in some systems, the TnsA endonuclease. The CASTs characterized to date differ primarily in the organization of their DNA targeting and transposition modes^1,2,16–18^. While type I variants typically use a multi-subunit Cascade-TniQ complex for crRNA-guided target recognition together with TnsA-TnsB for cut-and-paste transposition^19–21^, type V-K systems rely on the Cas12k-TniQ-S15 complex for crR-NA-guided targeting^5^ and use standalone TnsB for copy-and-paste transposition, resulting in cointegrate products^22^. TnsC plays a pivotal role in all CASTs, acting as an adaptor that bridges the targeting module and the transposase enzymes to define the insertion site^23,24^.

The inherent programmability of CASTs has driven their development as genome-editing tools that enable site-specific, double-strand break (DSB)-free insertion of multi-kilobase DNA cargoes in bacteria^25–28^ and mammalian cells^7,8,29,30^, offering considerable potential for research and therapeutic applications. While type V-K CASTs generally exhibit low integration efficiencies in human cell lines^29,30^, limited specificity^2,26,27,31^ and heterogeneous insertion profiles^22,30^, type I-F systems achieve precise, simple insertions with substantially higher efficiencies^1,7,22,26^. In particular, the type I-F system from the *Pseudoalteromonas* Tn*7016* transposon (**Fig. 1a**) has emerged as a highly promising candidate for genome engineering applications, distinguished by its uniquely high integration activity in human cells^7,9^. Directed evolution and structure-guided engineering of *Pse*CAST components in *E. coli* resulted in the evoCAST system, which exhibits markedly enhanced integration efficiencies, reaching up to 10–30% in human cells, while maintaining precise, cointegrate-free insertion outcomes^8^. While structural studies of type V-K CAST systems have yielded mechanistic insights into various stages of transpososome assembly^5,6,32–36^, structural information on type I CASTs is currently limited to sub-complexes^9,19–21,37–42^, presenting a knowledge gap in the mechanistic understanding of the RNA-guided DNA transposition pathway. Specifically, the mechanism by which the RNA-guided targeting module induces the assembly and catalytic activation of the transposase components remains elusive.

**Figure 1.**
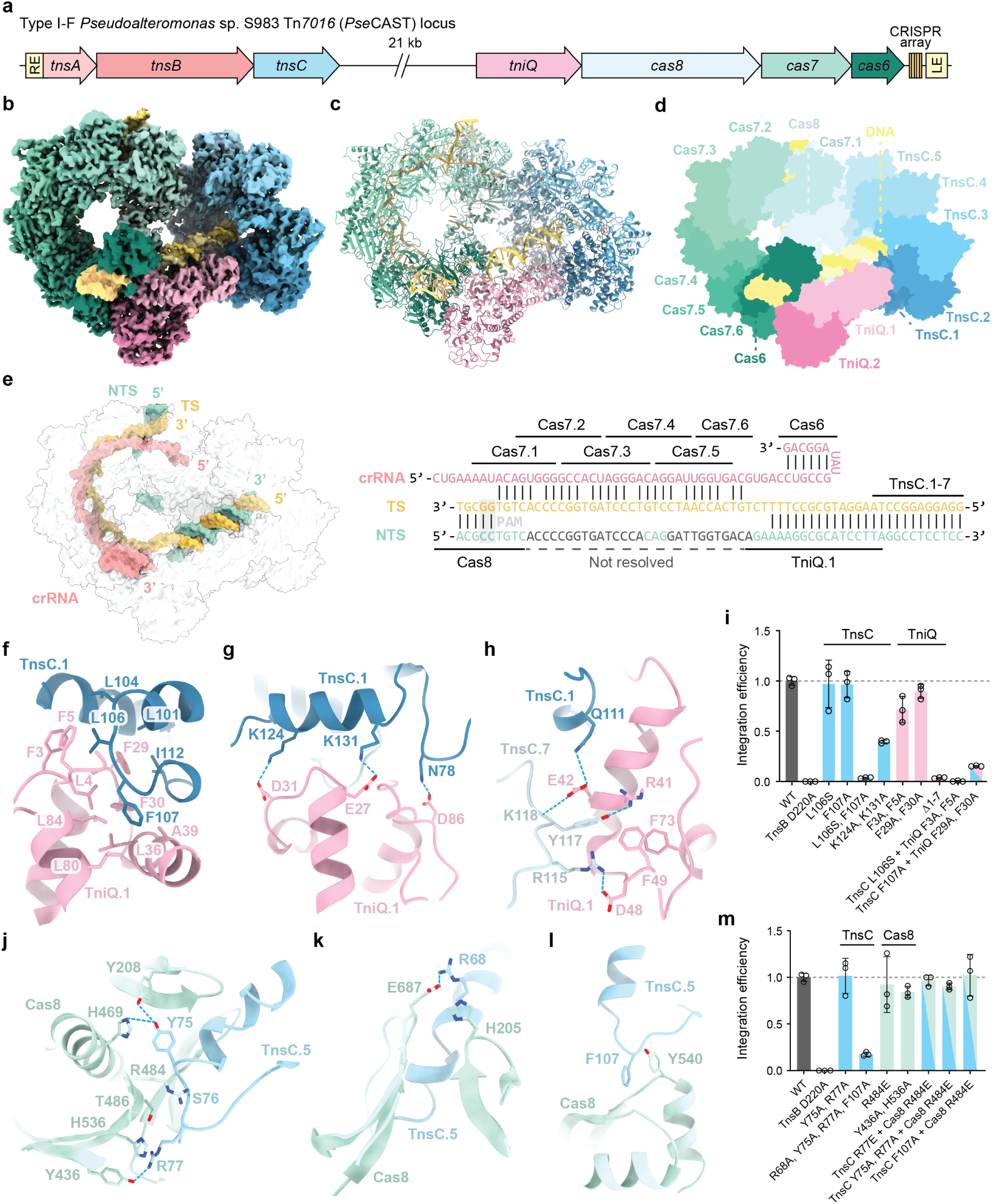
Molecular architecture of the *Pse*Cascade-TniQ-TnsC targeting complex. a, Locus organization of the type I-F CAST from *Pseudoalteromonas* sp. S983 (Tn*7016*). RE and LE denote transposon right and left ends, respectively. **b**, Cryo-EM density map of the *Pse*Cascade-TniQ-TnsC targeting complex bound to DNA, colored as in **a**. **c**, Atomic model of *Pse*Cascade-TniQ-TnsC, colored as in **b**. **d**, Schematic representation of the *Pse*Cascade-TniQ-TnsC complex, highlighting individual subunits. crRNA: CRISPR RNA. **e**, DNA substrate organization in the transposase recruitment complex (left), with an accompanying diagram summarizing strand identity and protein footprints derived from the model (right). TS, target strand; NTS, non-target strand. **f-h**, Detailed views of the interactions established between TniQ.1, TnsC.1, and TnsC.7. **i**, Normalized *E. coli* transposition activity of *Pse*CAST containing mutations in the TniQ-TnsC interface, as determined by qPCR. Data are presented as mean ± s.d. of three independent biological replicates (n = 3); the catalytically inactive TnsB mutant D220A serves as a negative control. **j-l**, Detailed views of the interaction interface between Cas8 and TnsC.5. **m**, Normalized *E. coli* transposition activity of *Pse*CAST containing mutations in the Cas8-TnsC interface, as determined by qPCR. Data are plotted as in **i**. transposition activity (Fig. 1i). Lastly, cross-subunit mutant combinations involving residues in the TnsC loop and the TniQ hydrophobic pocket (L106S^TnsC^ with F3A^TniQ^ and F5A^TniQ^; F107A^TnsC^ with F29A^TniQ^ and F30A^TniQ^) resulted in a marked decrease or the complete loss of transposition *in vivo* (Fig. 1i), confirming the importance of the TniQ-TnsC interactions for the assembly of the *Pse*CAST transpososome.

Here, we report a series of cryo-EM structural snapshots of the *Pse*CAST system, revealing the molecular determinants of RNA-guided DNA integration at key stages of the integration pathway, including the elusive transpososome holocomplex comprising all molecular components. The structure of the Cascade-TniQ-TnsC targeting complex reveals an unexpected molecular architecture in which interactions with both TniQ and Cas8 contribute to assembly of a TnsC heptamer on the target DNA. In turn, the structure of the *Pse*CAST transposase recruitment complex reveals that TnsC heptamerization is a prerequisite for the initial capture of the transposase, mediated by the C-terminal tail of TnsB. Finally, our structure of the 1.2 MDa *Pse*CAST holocomplex defines how the RNA-guided targeting machinery docks the TnsAB tetramer, along with the excised transposon donor DNA, onto the target DNA to ensure site-specific transposition. By trapping TnsAB in a strand-transfer state, this structure uncovers key TnsC-TnsB and TnsB-target DNA interactions that drive allosteric activation of the TnsB transposase to catalyze donor DNA integration. Collectively, our results elucidate the molecular mechanisms governing RNA-guided DNA integration in type I CASTs, unveiling the coordinated interplay between its modular components and establishing a structural foundation for its further development as a genome engineering technology.

## RESULTS

### Target-dependent assembly of TnsC heptamer by Cascade-TniQ

Our initial structural analysis by cryo-EM yielded a reconstruction of a target DNA-bound complex comprising *Pse*Cascade-TniQ and a heptameric TnsC ring, hereafter referred to as ‘targeting complex’, at an overall resolution of 3.1 Å (**Fig. 1b-d**, **Extended Data Fig. 1a-c**; **Extended Data Table 1**). Notably, the recruitment complex features a complete 32-bp R-loop (**Fig. 1e**), which was only partially formed in the previously reported intermediate *Pse*Cascade-TniQ complex structure^9^. In contrast to the open arrangement seen in the partial R-loop intermediate^9^, the Cas8 α-helical bundle (residues 272-421) undergoes a ∼90° rotation into a closed or locked state (**Extended Data Fig. 2a**), establishing additional contacts with the PAM-distal segment of the target DNA, as well as both TniQ subunits and Cas7.5 (**Extended Data Fig. 2b-d**). Moreover, three nucleotides of the displaced non-target strand (NTS) are bound in a pocket in the Cas8 bundle (**Extended Data Fig. 2e**), which likely facilitates NTS reannealing with the target strand (TS) in the PAM-distal DNA segment and its interactions with the TniQ dimer, a feature absent in partial R-loop and open helical-bundle states of type I-F CAST Cascade-TniQ complexes^19,21^. In line with our observations, the presence of the full R-loop and a closed Cas8 bundle conformation has been posited to represent a stable, target-engaged complex competent for transposase recruitment^21^. The formation of this locked state upon complete guide-target pairing suggests that it functions as a target fidelity checkpoint prior to transposase module recruitment.

**Figure 2.**
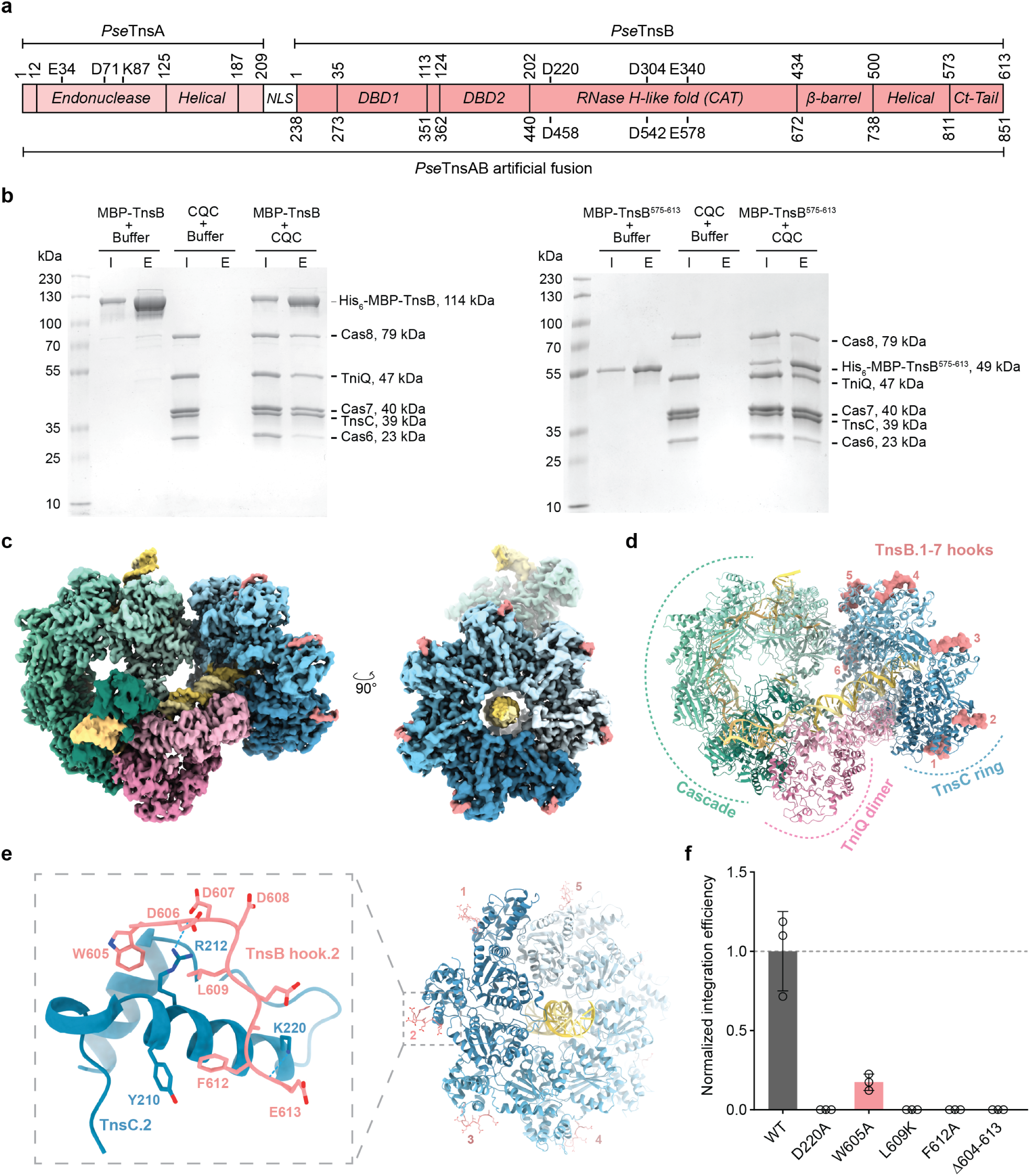
Recruitment of TnsB transposase via its C-terminal hook motif. a, Domain organization of *Pse*TnsA, *Pse*TnsB, and an artificial *Pse*TnsAB fusion protein. Numbers above the schematic refer to amino-acid positions for individual proteins; numbers below refer to the *Pse*TnsAB fusion protein. NLS, nuclear localization signal; DBD1/2, DNA-binding domain; CAT, catalytic domain; Ct-Tail, C-terminal tail. **b**, Co-precipitation of Cascade-TniQ-TnsC transposase recruitment complex by amylose-immobilized His_6_-MBP-tagged full-length TnsB (left) or C-terminal tail (right) in the presence of ATP and target dsDNA. CQC, Cascade-TniQ-TnsC; I, 5% input control; E, elution. **c**, Cryo-EM density map of the *Pse*Cascade-TniQ-TnsC transposase recruitment complex bound to TnsB-hook motifs and **d**, corresponding atomic model. **e**, Right: atomic model of the TnsB hook-decorated TnsC heptamer. Left: zoom-in view of the interactions established between TnsC.2 and the TnsB.2 hook motif. **f,** Normalized *E. coli* transposition activity of *Pse*CAST containing mutations in the TnsB hook, as determined by qPCR. Data are presented as mean ± s.d. of three independent biological replicates (n = 3); the catalytically inactive TnsB mutant D220A serves as a negative control.

The heptameric TnsC ring is assembled around the dsDNA exiting the Cascade-TniQ targeting complex (**Fig. 1d**), mirroring the architecture of TnsC heptamers from the canonical *Eco*Tn7 element and other type I CAST systems^20,40,42,43^ (**Extended Data Fig. 3**).The TnsC heptamer exhibits a near-perfect sevenfold symmetry with no helical rise or seam, fully enclosing the PAM-distal DNA duplex in a positively charged central channel, and all seven TnsC ATPase sites are occupied with ATP molecules and Mg^2+^ ions. The enclosed DNA adopts B-form geometry and is contacted by the Glu105-Asp120^TnsC^ loops of TnsC.1 and TnsC.5 via nonspecific interactions.

TnsC oligomers function as bridging modules for the assembly of the transposition machinery in both type V-K and type I-B CASTs, communicating with the CRISPR targeting effector and the transposase via their distinct N- and C-terminal faces, respectively^6,20,36,41^ (**Extended Data Fig. 3**). The *Pse-* CAST targeting complex structure reveals, unexpectedly, that the TnsC heptamer interacts with both TniQ.1 and Cas8 through extensive interfaces spanning 947 Å^2^ and 818 Å^2^, respectively (**Fig. 1d**; **Extended Data Fig. 3**), forming a ‘super-ring’ assembly that bridges the PAM-proximal and PAM-distal ends of the Cascade-TniQ subcomplex. At the N-terminal face of the heptamer, TnsC protomers use the same surfaces for interactions with both TniQ and Cas8 (**Extended Data Fig. 2f**). At the PAM-distal end, the N-terminal region of TniQ.1 is wedged into the TnsC.1-TnsC.7 inter-subunit interface, presenting an aromatic pocket comprising Phe3^TniQ^, Phe5^TniQ^, Phe29^TniQ^ and Phe30^TniQ^ to interact with a hydrophobic loop in TnsC.1 containing Leu101^TnsC^, Leu104^TnsC^, Leu106^TnsC^, Phe107^TnsC^ and Ile112^TnsC^ (**Fig. 1f**). TnsC.1 furthermore engages TniQ.1 in a series of ionic interactions (**Fig. 1g**), as well as bridging interactions that also contact the TnsC.1–TnsC.7 interface (**Fig. 1h**). Individual mutations within the inserted TnsC loop (L106S, F107A) did not impact the *E. coli* transposition activity of *Pse*CAST, while the double mutant exhibited a strong synergistic effect leading to a near-complete loss of transposition (**Fig. 1i**). While alanine substitutions of residues within the TniQ hydrophobic pocket (F3Aand F5A; F29A and F30A) had a mild effect on transposition efficiency, deletion of the seven N-terminal TniQ residues resulted in the loss of

**Figure 3.**
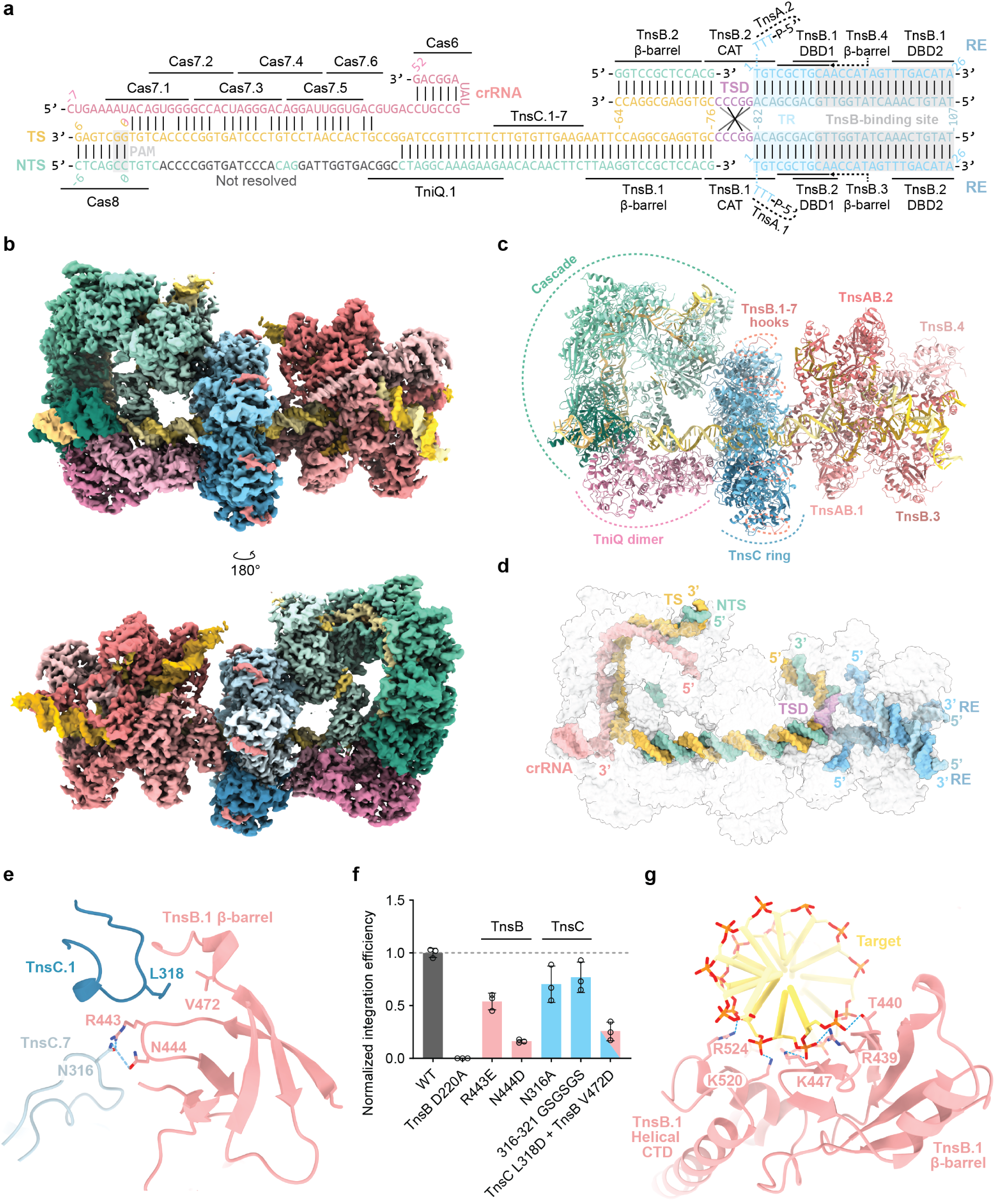
Molecular architecture of the 1.2 MDa *Pse*Cascade-TniQ-TnsC-TnsAB holocomplex. **a**, Schematic diagram of the strand-transfer DNA product mimic used for complex reconstitution, as modelled in the structure, with protein footprints highlighted. TS, target strand; NTS, non-target strand; crRNA, CRISPR RNA; TSD, target site duplication; TR, terminal repeat; RE, transposon right end. **b**, Cryo-EM density composite map of the *Pse*Cascade-TniQ-TnsC-TnsAB holocomplex. **c**, Atomic model of the holocomplex, with protein subunits coloured as in Fig. 2c; individual TnsAB subunits are depicted in varying shades of orange. **d**, Atomic model of the strand-transfer DNA product, with distinct regions highlighted in different colours. **e**, Detailed view of the interactions of the beta-barrel domain of TnsB.1 with TnsC.1 and TnsC.1/7 subunits. **f**, Normalized *E. coli* transposition activity of *Pse*CAST containing mutations in the TnsC-TnsB beta-barrel domain interface, as quantified by qPCR. Data are presented as mean ± s.d. of three biologically independent replicates (n = 3). **g**, Overview of the beta-barrel and C-terminal helical TnsB domains contacting the target DNA duplex.

At the PAM-proximal end, Cas8 and TnsC.5 form a shape-complementary interface, mediated by aromatic and electrostatic interactions (**Fig. 1j–l**). Combined alanine substitutions of Arg68^TnsC^, Tyr75^TnsC^, Arg77^TnsC^, and Phe107^TnsC^ resulted in a pronounced reduction of transposition *in vivo* (**Fig. 1m**), indicating that the Cas8-TnsC interaction contributes to transpososome assembly. However, mutations of interacting residues in Cas8, or their cross-subunit combinations, had a mild or negligible effect (**Fig. 1m**), suggesting a lower sensitivity of the TnsC-Cas8 interface to mutational perturbations as compared to the TnsC-TniQ interface. Together, these structural observations define the molecular coupling between the CRISPR targeting machinery and TnsC that results in the assembly of an ATP-loaded TnsC heptamer on the target DNA.

### Initial transposase recruitment via C-terminal tail of TnsB

In the canonical *E. coli* Tn7 element^44^, as well as in *Sh*CAST (type V-K) and *Pmc*CAST (type I-B) systems, transposase recruitment is mediated by TnsC interactions with C-terminal regions of TnsB^6,23,41^. In the structure of the *Pse*TnsAB paired-end complex (PEC), the C-terminal tail of *Pse*TnsB (residues 574-613) is disordered^45^. To assess whether the unstructured tail plays a similar role in engaging *Pse*TnsC, we performed co-precipitation experiments using affinity-tagged *Pse*TnsB constructs comprising either the full-length protein or the standalone C-terminal tail (TnsB^575–613^) (**Fig. 2a**). Both full-length TnsB and TnsB^575–613^ efficiently co-precipitated target DNA-bound Cascade-TniQ-TnsC in the presence of ATP (**Fig. 2b**), indicating that TnsB directly interacts with the transposase recruitment complex and that its C-terminal tail is sufficient to mediate this interaction. Furthermore, the amount of co-precipitated TnsC remained unchanged in the presence of the non-hydrolysable ATP analogue ATPγS (**Extended Data Fig. 4a**), implying that TnsB does not stimulate ATPase-dependent depolymerisation of TnsC in *Pse*CAST, similar to the type I-B *Pmc*CAST system^41,42^.

To validate these findings structurally, we aimed to reconstitute and determine the structure of Cascade-TniQ-TnsC in complex with TnsAB bound to the excised transposon ends. To this end, we performed affinity co-precipitation using a recombinant TnsAB fusion protein construct (**Fig. 2a**) pre-assembled with a fragment of the transposon right end (RE), to capture the Cascade-TniQ-TnsC complex assembled on a linear dsDNA target substrate (**Extended Data Fig. 4b**). Single-particle cryo-EM analysis of the resulting sample yielded a 2.8 Å reconstruction of the *Pse*CAST transposase recruitment complex comprising the target-bound Cascade-TniQ-TnsC and additional densities corresponding to the TnsB C-terminal segments (residues Trp605-Glu613), hereafter referred to as ‘TnsB hooks’. The TnsB hooks are bound in all TnsC subunits at the junction between the N-terminal α/β core domain and the C-terminal α-helical bundle in proximity to the ATP binding pocket (**Fig. 2c,d**; **Extended Data Fig. 5a-c**; **Extended Data Table 1**), similar to type V-K^34,36^ and I-B CASTs^41,42^ (**Extended Data Fig. 6**). The TnsB hook is anchored by hydrophobic interactions of Trp605^TnsB^, Leu609^TnsB^ and Phe612^TnsB^, and additional hydrogen bonding between the backbone carbonyl of Asp606^TnsB^ and Arg212^TnsC^ (**Fig. 2e**). Corroborating these observations, alanine substitution of Trp605^TnsB^ resulted in a substantial reduction of *E. coli* transposition activity of *Pse*CAST, while the individual substitutions L609K^TnsB^ and F612A^TnsB^, or deletion of the entire TnsB hook (Δ604-613), completely abolished transposition (**Fig. 2f**), highlighting the functional requirement of TnsB hook-TnsC interactions for *Pse*CAST transposition. These findings demonstrate the critical role of the TnsB C-terminal tail for the recruitment of the TnsAB transposase and delivery of the transposon DNA to the target site prior to integration.

### Transpososome holocomplex assembly

We next reconstituted a holocomplex comprising Cascade, TniQ, TnsC, and TnsAB by a ‘one-pot’ affinity co-precipitation approach similar to the one used for the assembly of the TnsB hook-bound recruitment complex. A symmetric DNA substrate mimicking the strand-transfer intermediate was used to trap the transpososome in a post-integration, strand-transfer complex (STC) state (**Fig. 3a**, **Extended Data Fig. 7**), whose stability has been previously leveraged to visualize transpososome structures^6,41,46^. Cryo-EM analysis of the resulting sample, aided by 3D variability analysis and heterogeneous and locally-focused refinements, yielded a reconstruction with a global resolution of 3.1 Å (**Fig. 3b**, **Extended Data Fig. 8a-d**; **Extended Data Table 1**). The reconstruction reveals that the transposase adopts an intertwined, pseudo-twofold arrangement with a TnsA_2_-TnsB_4_ stoichiometry to assemble on the structured DNA in a STC configuration (**Fig. 3c,d**). Additionally, all seven TnsC subunits are observed bound to the C-terminal hook motifs of TnsAB (**Fig. 3b,c**), although it is not possible to ascertain which of these are connected to the bound TnsAB molecules, as the connecting segments (residues 824-843) are unstructured. The target DNA exits from the central channel of the TnsC heptamer and extends towards the TnsAB tetramer assembled on the pseudo-palindromic target site duplication (TSD) region and the adjacent transposon end sequences (**Fig. 3a**). As in the structure of the TnsAB paired-end complex^45^, each transposon end is occupied by two TnsB protomers: a proximal ‘catalytic’ subunit (TnsB.1/2) whose catalytic domain engages the other transposon end in *trans*, and a distal ‘structural’ subunit (TnsB.3/4) (**Fig. 3c**). Only two TnsA subunits (TnsA.1/2), bound to the catalytic TnsB protomers, are resolved in the structure. The corresponding interaction interfaces in the structural TnsB subunits are occupied by the bound RE DNA duplexes, precluding TnsA interactions.

The structure of the *Pse*CAST holocomplex defines molecular interactions responsible for Tn-sC-dependent recruitment of TnsAB to the target DNA to effect site-specific donor integration. The TnsB.1 subunit is tethered to the C-terminal face of the TnsC heptamer by its beta-barrel domain (residues 434-500), positioned at the interface of TnsC.1 and TnsC.7 subunits (**Fig. 3e**). TnsC.1 inserts its C-terminal loop spanning residues 314-322, including Leu318^TnsC^, into a hydrophobic pocket in the TnsB.1 beta-barrel domain, lined with Val472, Gln467, and Leu441^TnsB^ (**Fig. 3e**). In turn, Asn316^TnsC^, situated in the same loop in TnsC.7, engages in hydrogen-bonding interactions with Arg443^TnsB^ and Asn444^TnsB^ (**Fig. 3e**). Mutating both Leu318^TnsC^ and Val472^TnsB^ to aspartate resulted in a pronounced reduction of transposition *in vivo* (**Fig. 3f**). Although a single alanine substitution of Asn316^TnsC^ as well as replacement of the TnsC loop residues 316-321 with a (Gly-Ser)_3_ linker resulted in modestly impaired activity, single mutations of Arg443^TnsB^ or Asn444^TnsB^ strongly reduced transposition (**Fig. 3f**), confirming that formation of this interface is essential for *Pse*CAST-mediated DNA transposition.

In addition to interactions with TnsC, the beta-barrel domain of the catalytic TnsB.1 subunit also interacts with the backbone of the target DNA segment downstream of the TnsC heptamer, forming a network of hydrogen bonding and electrostatic contacts using Arg439^TnsB^, Thr440^TnsB^, Lys447^TnsB^, Gln449^TnsB^, and Glu450^TnsB^ (**Fig. 3g**). Furthermore, Lys520^TnsB^ and Arg524^TnsB^ in the C-terminal helical domain also contact the target DNA backbone. Notably, this domain adopts a helical bundle fold that bridges the catalytic and beta-barrel domains in the catalytic TnsB.1/2 subunits, in contrast to the structural TnsB.3/4 sub-units, in which the helical domain forms two stretched helices that are sandwiched between the catalytic and DBD2 domains of TnsAB.1/2, contributing to intra-subunit contacts.

Taken together, these insights highlight the functional role of the TnsB beta-barrel domain in transpososome assembly. The shape complementarity of the TnsC and TnsB beta-barrel interaction surfaces, together with cooperative interactions with the target DNA, provide the structural basis for the molecular ruler mechanism underpinning integration site definition, facilitating transposon insertion at a fixed distance of 49 nt from the crRNA-specified target site. Moreover, the synergistic contacts with two TnsC subunits and the target DNA underlie a checkpoint mechanism that senses target DNA-dependent TnsC oligomerization, thereby explaining the strictly sequential transpososome assembly observed in type I-F systems^40^.

### Target-dependent transposase activation

The *Pse*CAST transpososome holocomplex structure captures the TnsAB STC tetramer in a post-integration state (**Fig. 4a,b**). As in the PEC structure^45^, the active sites of the TnsA.1/2 subunits engage the 5’ termini of the transposon ends, adopting a catalytically competent, post-cleavage geometry in which the catalytic triad residues Glu34^TnsA^, Asp71^TnsA^, Lys87^TnsA^ coordinate the 5′-terminal phosphate group and a single Mg²^+^ ion (**Fig. 4c**). In contrast to the PEC structure, the catalytic sites of TnsB.1/2 subunits also assume an activated conformation in which a Mg²^+^ ion, coordinated by the DDE catalytic triad Asp220^TnsB^, Asp304^TnsB^, and Glu340^TnsB^, interacts with the 3′-hydroxyl group of the terminal nucleotide (dG76 in the NTS) of the target DNA and the phosphodiester group linking the TSD segment with the 3’-terminal nucleotide of the transposon end (**Fig. 4d**). In addition, the catalytic domain establishes contacts with RE and TSD nucleotides that are critical for STC formation (**Fig. 4d**). Specifically, Asp225^TnsB^ positions an additional Mg^2+^ ion to simultaneously coordinate the phosphate groups of dA-82 and dG-80, enabling a kink in the DNA backbone at the TSD-RE junction to facilitate STC formation. The backbone is additionally contacted by Thr222^TnsB^, Arg239^TnsB^, and Ser360^TnsB^ (**Fig. 4d**). In agreement with these observations, alanine substitution of Asp225^TnsB^ or charge reversal mutation of Arg239^TnsB^ resulted in complete loss of transposition activity *of Pse*CAST in *E. coli*, while mutations of Thr222^TnsB^ or Ser360^TnsB^ reduced activity (**Fig. 4e**).

**Figure 4.**
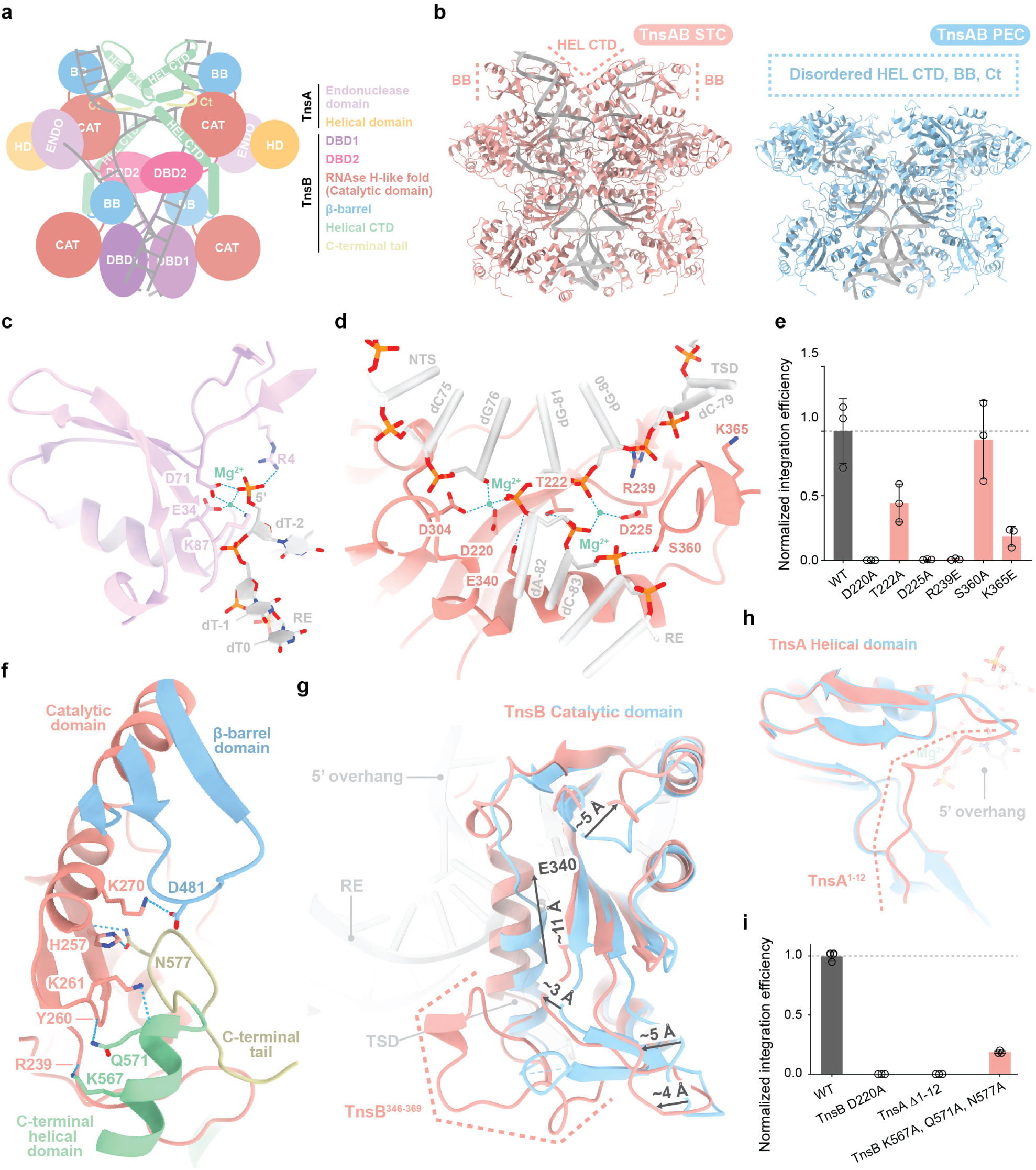
TnsC- and target DNA-dependent allosteric activation of TnsB. **a**, Schematic diagram of the TnsAB strand-transfer complex (STC), with individual domains coloured according to the figure legend. Each domain is labelled with the corresponding domain-name abbreviation. **b**, Structures of TnsAB bound to DNA (grey) in the STC state within the *Pse*CAST holocomplex (left) and in the paired-end complex (PEC, right) conformations. TnsB domains that become structured in the STC (BB, beta-barrel domain; HEL CTD, helical C-terminal domain; Ct, C-terminal tail) are indicated. **c**, Zoom-in view of the 5’-terminal transposon end overhang in the catalytic site of TnsA. RE, transposon right end. **d**, Zoom-in view of the TnsB catalytic site and interactions between the target site duplication (TSD) DNA and the TnsB catalytic domain. NTS, non-target strand. **e**, RNA-guided transposition activity in *E. coli* for *Pse*CAST systems containing mutations in the TnsB residues involved in contacting the TSD region of target DNA, as quantified by qPCR. Data are presented as mean ± s.d. of three biologically independent replicates (n = 3). Black circles represent individual measurements. **f**, Detailed view of the interface between TnsB beta-barrel, C-terminal helical, C-terminal tail, and catalytic domains in the PTC. **g,h,** Zoom-in views of the structural overlays of TnsAB in the PEC (blue) and STC (orange) states, highlighting (**g**) allosteric remodeling of the TnsB catalytic domain and ordering of the 346-369 region, shift in the helix carrying the catalytic residue Glu340^TnsB^, and (**h**) ordering of the first 12 N-terminal residues of TnsA. **i**, RNA-guided transposition activity in *E. coli* of *Pse*CAST systems containing mutations in the N-terminus of TnsA and in the residues at the interface shown in **f**, as quantified by qPCR. Data are presented as mean ± s.d. of three biologically independent replicates (n = 3). Black circles represent individual measurements.

Finally, the *Pse*CAST transpososome holocomplex structure illuminates a conformational relay mechanism underlying target DNA-dependent catalytic activation of TnsB. In the PEC structure, the beta-barrel and helical domains of the catalytic TnsB subunits are disordered^45^. By contrast, both domains are structured in the STC state and interface with the catalytic domain (**Fig. 4b**). This is achieved by a salt bridge between Asp481^TnsB^-Lys270^TnsB^ and a network of hydrogen bonding and ionic interactions formed between the catalytic domain and the C-terminal tail residues (Lys567^TnsB^-Val585^TnsB^) (**Fig. 4f**). This is associated with a remodelling of the catalytic domain, resulting in the ordering of an adjacent loop (residues Glu348-Lys369) (**Fig. 4g**) and a concomitant ∼11 Å shift of the helix bearing the catalytic glutamate Glu340^TnsB^ (**Fig. 4g**). The conformational rearrangement marks the transition of the catalytic site from an inactive, post-excision state to an active, strand-transfer configuration supporting Mg^2+^ coordination and catalytic activity. In agreement with these observations, combined alanine substitutions of Lys567^TnsB^, Gln571^TnsB^, and Asn577^TnsB^ strongly reduced the *E. coli* transposition activity of *Pse*CAST. Additional ordering is observed for the N-terminal segment (residues Met1-Pro11) of the TnsA helical domain, which positions Arg4^TnsA^ to contact the 5′-terminal phosphate of the cleaved transposon end (**Fig. 4c,h**), implicating TnsA in TnsB activation. Accordingly, truncation of the TnsA N-terminal segment (ΔMet1-Asn12^TnsA^) resulted in complete loss of transposition (**Fig. 4i**). Taken together, the structural observations and their functional validation indicate that the molecular interfaces formed between the C-terminal, beta-barrel, and catalytic domains of TnsB formed upon target DNA and TnsC binding are required for the integration activity of *Pse*CAST. In this way, both target DNA and TnsC synergistically couple transposase recruitment with domain ordering and catalytic site remodelling, resulting in allosteric activation of TnsB for site-specific DNA integration.

## DISCUSSION

Our structural and functional analysis of the *Pse*CAST system comprehensively reveals the molecular logic of type I-F RNA-guided transposition, in which an orchestrated series of interactions and conformational rearrangements couples target selection to achieve cut-and-paste DNA integration. Together, these insights support a mechanistic model in which type I-F CAST transposition is controlled by a succession of molecular checkpoints (**Fig. 5**).

**Figure 5.**
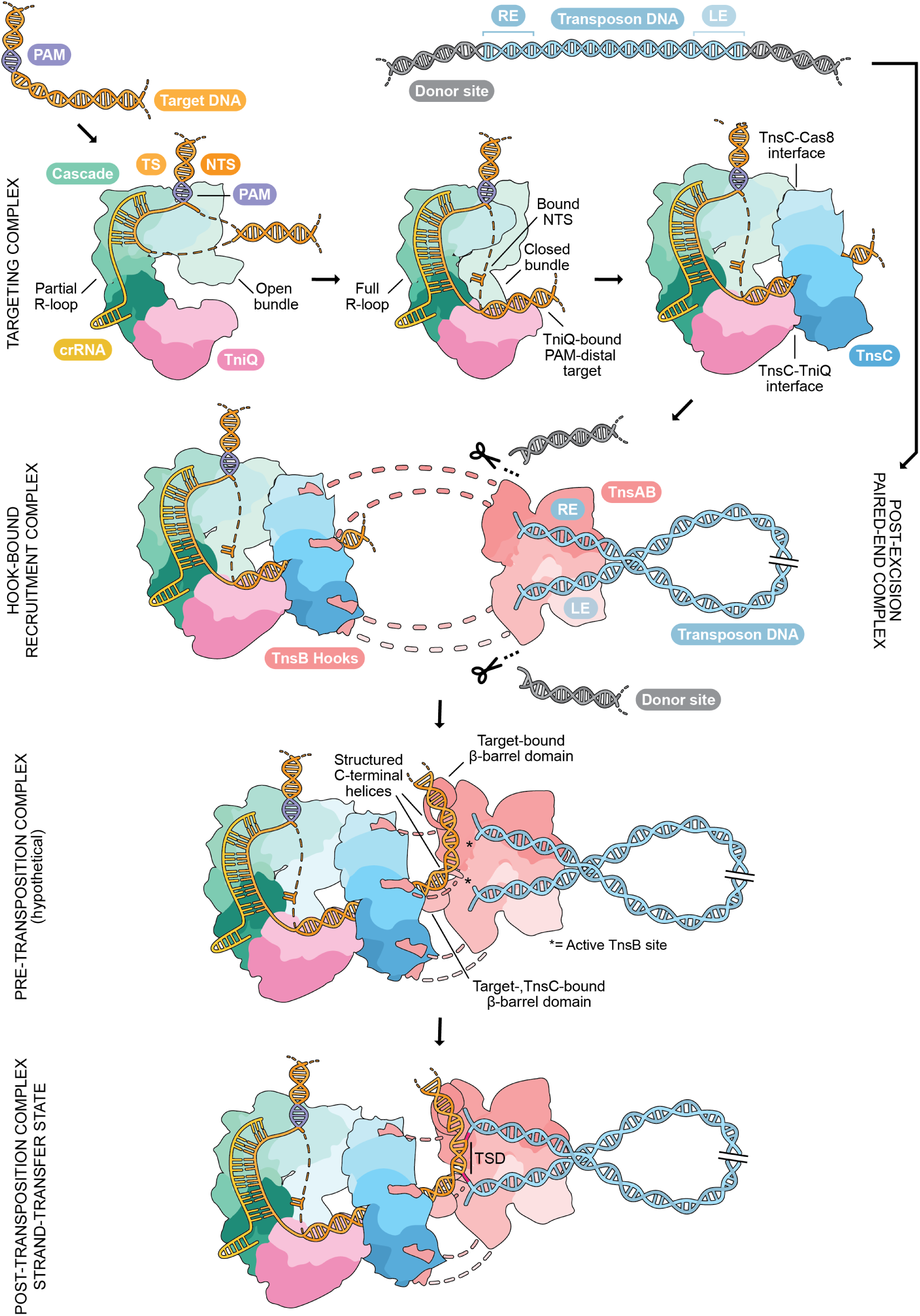
Mechanistic model of RNA-guided DNA-integration in type I-F CASTs. The Cascade-TniQ complex is guided to the target DNA by base-pairing with its crRNA, with initial formation of a partial R-loop. Coordination of the displaced NTS by the Cas8 helical bundle is coupled with its transition from open to closed, full R-loop formation, and stabilization of the double-stranded PAM-distal target by the TniQ dimer. TnsC is recruited at the target site, where it assembles in the presence of ATP into a closed heptameric ring by binding the PAM-distal target duplex in the presence of ATP, TniQ, and Cas8. At the donor site, cleavage of the transposon DNA by TnsAB proceeds through the formation of a paired-end complex. The PEC is initially recruited by the TnsC heptamer by interactions with the C-terminal TnsB hooks. Interactions with the target DNA and TnsC result in remodelling of the beta-barrel and C-terminal helical domains of the interfacing TnsB copies, which allosterically induces a rearrangement of the catalytic domains into an active conformation. The transposon DNA is integrated by two transesterification reactions catalyzed by the active sites of the interfacing TnsBs, connecting both transposon 3′-ends to the target DNA with 5-bp spacing that result in a target-site duplication (TSD).

The formation of the Cascade-TniQ-TnsC transposase recruitment complex requires R-loop completion, enabled by a conformational rearrangement of the Cas8 subunit whose helical bundle undergoes a ∼90° rotation to adopt a locked conformation. In the locked state, the helical bundle stabilizes the displaced non-target DNA strand and mediates its reannealing at the PAM-distal end of the R-loop. In contrast to open conformations observed in previously reported type I-F CAST Cascade-TniQ complexes^9,19,21,37^, this locked state presents a double-stranded PAM-distal target DNA for binding TniQ, which in turn facilitates ATP-dependent TnsC oligomerization. As this conformation is only achieved with fully matched targets, it acts as a fidelity checkpoint that discriminates against mismatched sites prior to TnsC recruitment. The architecture of the *Pse*CAST transposase recruitment complex highlights mechanistic differences from type V CASTs in coupling the CRISPR effector module to the transposon machinery. Instead of forming extended filaments that remodel DNA^32,33^, *Pse*TnsC assembles into an ATP-bound, symmetric heptamer that fully encircles the duplex exiting the TniQ dimer. In doing so, TnsC makes additional contacts with Cas8, forming a closed super-ring that clamps the targeting complex and fixes the orientation of the downstream DNA. The spacing between the Cascade R-loop and the insertion site is thus determined in part by the footprint and register of the Cascade-TniQ-TnsC recruitment complex, in contrast to TnsB-dependent TnsC depolymerization found in type V-K systems^6,32,33^.

The paired-end TnsAB transposase complex is initially flexibly tethered to the TnsC oligomer via the short C-terminal tail hook motifs of TnsB, and productive target DNA engagement requires the formation of an additional key interface between the TnsC heptamer and the beta-barrel domain of the catalytic TnsB subunit. Upon engagement of the target DNA, the beta-barrel domain and C-terminal helices of the TnsC-proximal catalytic TnsB subunits are positioned to interact with the catalytic domain. This conformational rearrangement is finally coupled to remodelling of the TnsB active site into a catalytically competent configuration by an allosteric relay mechanism. In this way, the TnsC- and target DNA-induced catalytic activation of TnsB acts as an additional fidelity checkpoint, ensuring that transposon integration proceeds only once the transposase is correctly anchored on the target DNA.

Comparison with other subtypes highlights both conserved principles and subtype-specific solutions by which CASTs use AAA+ ATPase oligomeric platforms to connect RNA-guided targeting modules to the transposon machinery. In type V-K systems, the analogous beta-barrel domain of TnsB also contacts TnsC in the strand-transfer complex and is required to stimulate TnsC ATPase-dependent filament disassembly^6^. In the type I-B *Pmc*TnsABCD complex the beta-barrel domain is positioned near the TnsC ring to form additional contacts^41^, but its functional relevance is not yet clear. In all three systems, the C-terminal helices fold back and interact with the TnsB catalytic domain^6,41^, whereas they are disordered in the

TnsC-less type V-K *Sh*TnsB complex^34,35^, analogously to what is observed for the type *Pse*TnsAB PEC. Overall, TnsB recruitment through a short C-terminal tail and regulation by peripheral domains is a shared feature, while the distinct TnsC architectures dictate distinct mechanisms of coupling target recognition to transposon recruitment, ATPase activation, and integration site selection. For type I CASTs, the position of the integration site is primarily defined by the architecture of the Cascade-TniQ-TnsC recruitment complex, its interactions with the TnsB beta-barrel domain, and organization of TnsAB at the transposon ends. Our structural data reveal an allosteric network linking target DNA engagement to transposase activation, but the sequence of interactions and activation steps may involve additional intermediates. Taken together, our findings illuminate the molecular mechanisms underpinning RNA-guided transposition in type I CASTs, providing a structural framework for their further rational engineering for technological applications.

## METHODS

### Plasmid construction

The expression plasmids containing His_10_-TEV-tagged TniQ with Cas8 (pSL4521), Cas7 with Cas6 and the CRISPR array (pSL4522), His_6_-SUMO-TEV-tagged TnsC (pSL5536), and His_6_-SUMO-TEV-tagged TnsAB fusion protein (pSL3768) were cloned using standard molecular biology methods including Gibson^47^ and Golden Gate assembly^48^. All plasmids were sequence-verified via whole-plasmid nanopore sequencing (Genewiz). For cloning of the expression plasmid containing Cas7, the CRISPR array and Twin-Strep-tagged Cas6 (pMS321), a Twin-Strep-tag^49^ was added at the C-terminus of the *cas6* gene by Gibson assembly: pSL4522 was linearized by PCR using oMS603 and oMS604, and the Twin-Strep-tag was amplified by PCR from a common backbone with primers oMS605 and oMS606, bearing compatible overhangs to the linearized pSL4522. For cloning of MBP-tagged TnsB and TnsAB fusion proteins, a His_6_-MBP-tagged version of TnsAB fusion and TnsB was generated by PCR amplification of the respective gene using pSL3768 as the template and the primers oMS631-632 and oMS634-632, respectively. The PCR product was inserted in the 1C backbone (Addgene #29654) by ligation-independent cloning^50^ (LIC). The His -MBP-tagged TnsB C-terminal tail (residues 575 to 613) was prepared by Gibson assembly, generating the PCR fragments using the His_6_-MBP-TnsB plasmid as a template and the primers pairs oGF181-182 and oGF180-184. Plasmids were cloned and propagated in Mach I cells (Thermo Fisher Scientific). Plasmids were purified using the GeneJET plasmid miniprep kit (Thermo Fisher Scientific) and verified by full-plasmid sequencing or Sanger sequencing (Microsynth). All sequences of primers and parental plasmids used for cloning are provided in **Supplementary Tables 1 & 2**.

### Protein expression and purification

For Cascade-TniQ protein expression, the plasmid pSL4521, encoding His_10_-TEV-TniQ and Cas8, and the plasmid pMS321, encoding Cas8, Cas6-Twin-Strep and the crRNA, were co-transformed in 50 μL of *E. coli* BL21 Rosetta2 (DE3) cells (Novagen), plated on an agar plate and incubated at 37 °C overnight. A single colony was inoculated in 100 mL of Luria Broth (LB) to obtain a starting culture. Both transformation plate and liquid media were supplemented with 50 µg/mL kanamycin, 50 µg/mL streptomycin and 33 µg/mL chloramphenicol. After overnight growth at 37 °C, 220 rpm, the starting culture was added to eight antibiotic-supplemented (kanamycin, streptomycin) LB media flasks, each 750 mL, to a starting optical density at 600 nm (OD_600_) of 0.05. Cell cultures were grown at 37 °C, shaking at 110 rpm, until an OD_600_ of 0.6 was reached. Protein expression was induced with 0.4 mM IPTG and continued for 16 h at 18 °C. Cells were collected by centrifugation (4,000*g* for 30 min at 4 °C) and resuspended in 100 mL lysis buffer (20 mM Tris-HCl pH 8.0, 500 mM NaCl, 5 mM imidazole) supplemented with 1 μg/mL pepstatin and 200 μg/mL AEBSF. The suspension was lysed using a Maximator cell homogenizer at 1,500 bar and 4 °C. The lysate was cleared by centrifugation (40,000*g* for 40 min at 4 °C) and applied to three 5-mL HisTrap HP columns (Cytiva) connected in tandem and equilibrated in lysis buffer. The columns were washed with 200 mL of HisTrap wash buffer (20 mM Tris-HCl pH 8.0, 500 mM NaCl, 5 mM imidazole) and eluted with 50 mL of HisTrap elution buffer (20 mM Tris-HCl pH 8.0, 500 mM NaCl, 250 mM imidazole). DTT and ATP were added to the elution to a final concentration of 1 mM each. The sample was added to 7 mL of 50% Strep-Tactin affinity resin suspension (IBA Lifesciences) in HisTrap elution buffer. The mixture was incubated for 30 min at 4 °C on a rotating wheel. The protein-bound resin was washed with 70 mL of size exclusion (SEC) buffer (20 mM HEPES-KOH pH 7.5, 500 mM KCl, 1 mM DTT) and the sample was eluted with 20 mL of SEC buffer supplemented with 5 mM D-Desthiobiotin. Eluted fractions were further purified by SEC using a Superose 6 Increase 10/300 GL (Cytiva) column in SEC buffer. Purified proteins were concentrated to ∼10 mg/mL using a 30 kDa molecular weight cut-off centrifugal filter (Merck Millipore), flash frozen in liquid nitrogen and stored at −80°C until further use.

For TnsC expression, vector pSL5536, encoding His_6_-SUMO-TEV-TnsC, was transformed in 50 μL of *E. coli* BL21 Star (DE3) cells (Thermo Fisher Scientific), plated on an agar plate supplemented with 50 µg/mL kanamycin and incubated at 37 °C overnight. A single colony was inoculated in 100 mL of kanamycin-supplemented LB to obtain a starting culture. After overnight growth at 37 °C, 220 rpm, the starting culture was added to eight kanamycin-supplemented LB media flasks, each 750 mL, to a starting OD_600_ of 0.05. Cell cultures were grown at 37 °C shaking at 110 rpm until an OD_600_ of 0.6 was reached.

Protein expression was induced with 0.3 mM IPTG and continued for 16 h at 18 °C. Cells were collected by centrifugation (4,000*g* for 30 min at 4 °C). The purification followed the protocol published for *Vch*TnsC in Hoffman *et al.* ^40^: a cell pellet (corresponding to four 750 mL liquid cultures) was resuspended in 40 mL of lysis buffer (20 mM HEPES-KOH pH7.5, 1 M NaCl, 5 mM magnesium acetate, 0.2 mM EDTA, 0.1% (v/v) Triton X-100, 5% (v/v) glycerol, 1 mM DTT) supplemented with 1 μg/mL pepstatin, 200 μg/ mL AEBSF, and 1 μg/mL DNAse. The suspension was lysed by ultrasonication (Bandelin Sonopuls HD 3200 equipped with a VS 70 T probe, 1 s /2 s on/off cycle, at 20% amplitude, for 15 min, at 4 °C). The lysate was cleared by centrifugation (40,000*g* for 45 min at 4 °C) and applied to two 5-mL HisTrap HP columns (Cytiva) connected in tandem and equilibrated in equilibration buffer (20 mM HEPES-KOH pH 7.5, 1 M NaCl, 0.2 mM EDTA, 5% (v/v) glycerol, 10 mM imidazole, 1 mM DTT). The columns were washed with 90 mL of equilibration buffer and with 50 mL of wash buffer (20 mM HEPES-KOH pH 7.5, 1 M NaCl, 0.2 mM EDTA, 5% (v/v) glycerol, 30 mM imidazole, 1 mM DTT), and the proteins were eluted with 10 mL of elution buffer (20 mM HEPES-KOH pH 7.5, 1 M NaCl, 0.2 mM EDTA, 5% (v/v) glycerol, 300 mM imidazole, 1 mM DTT). Elution fractions were pooled and dialyzed overnight against 2 L of dialysis buffer (20 mM HEPES-KOH pH 7.5, 1 M NaCl, 0.2 mM EDTA, 1% (v/v) glycerol, 1 mM DTT, 10 mM imidazole) in the presence of 2% (w/w) His_6_-tagged TEV protease. Any precipitated protein was removed by centrifugation (20,000*g* for 20 min at 4 °C). The dialyzed proteins were applied to two 5-mL HisTrap HP columns (Cytiva) connected in tandem in equilibration buffer. The columns were washed with 60 mL of equilibration buffer, 90 mL of wash buffer, and 50 mL of elution buffer. Flowthrough and initial wash fractions were applied to a Superdex 200 Increase 10/300 GL column (Cytiva) equilibrated in SEC buffer (20 mM HEPES-KOH pH 7.5, 1 M NaCl, 0.2 mM EDTA, 1% (v/v) glycerol, 1 mM DTT). Purified proteins were concentrated to ∼12 mg/mL using a 10 kDa molecular weight cut-off centrifugal filter (Merck Millipore), flash frozen in liquid nitrogen and stored at −80°C until further use.

For the expression of His_6_-SUMO-TEV-TnsAB (pSL3768), His_6_-MBP-TEV-TnsAB (pMS329), His_6_ -MBP-TEV-TnsB (pMS331) and His_6_ -MBP-TEV-TnsB^575–613^ (pGF189), the same procedure as for His_6_-SUMO-TEV-TnsC was followed, except that 0.4 mM IPTG was used to induce protein expression instead of 0.3 mM. For the purification of untagged TnsAB, TnsB, or TnsB^575-613^, a cell pellet (corresponding to four 750 mL liquid cultures) was resuspended in 40 mL of lysis buffer (20 mM Tris-HCl, pH 7.5, 500 mM NaCl, 5 mM imidazole, 5% glycerol) supplemented with with 1 μg/mL pepstatin, 200 μg/mL AEBSF, and 1 μg/mL DNAse. The suspension was lysed by ultrasonication (Bandelin Sonopuls HD 3200 equipped with a VS 70 T probe, 1 s/2 s on/off cycle, at 30% amplitude, for 15 min, at 4°C). The lysate was cleared by centrifugation (40,000*g* for 1 h at 4 °C) and applied to two 5-mL HisTrap HP columns (Cytiva) connected in tandem and equilibrated in lysis buffer. The columns were washed with 50 mL of lysis buffer, and the proteins were eluted with a stepwise increase (5%, 20% and 100%) of elution buffer (20 mM Tris-HCl, pH 7.5, 500 mM NaCl, 500 mM imidazole, 5% glycerol). Fractions corresponding to 20% and 100% of elution buffer percentage were pooled and dialyzed overnight against 2 L of dialysis buffer (20 mM Tris-HCl, pH 7.5, 500 mM NaCl, 5% glycerol, 1 mM DTT) in the presence of 2% (w/w) His_6_-tagged TEV protease. Dialyzed proteins were loaded in two 5-mL HiTrap Heparin HP columns (Cytiva) connected in tandem in dialysis buffer, washed with 30 mL of dialysis buffer and eluted with a linear gradient of heparin buffer (20 mM Tris-HCl, pH 7.5, 1 M NaCl, 5% glycerol, 1 mM DTT) over 150 mL (target concentration of heparin buffer: 100%). Peak fractions were pooled and further purified by size-exclusion chromatography using a Superdex 200 Increase 10/300 GL column (Cytiva) in SEC buffer (20 mM Tris-HCl, pH 7.5, 500 mM NaCl, 1 mM DTT). Purified proteins were concentrated to ∼15 mg/mL using a 50 kDa molecular weight cut-off centrifugal filter (Merck Millipore), flash frozen in liquid nitrogen and stored at −80°C until further use. For the expression and purification of His -MBP-tagged TnsAB/TnsB/TnsB^575-613^, the same procedure as for untagged TnsAB/TnsB/ TnsB^575-613^ was followed, with the only difference being that the TEV-cleavage and heparin steps were omitted.

### Pull-down co-precipitation experiments

A double-stranded target DNA (dsTarget) was prepared by annealing oligos oMS628 and oMS629 (**Supplementary Table 3**) in a 1:1 ratio to a final concentration of 100 μM in annealing buffer (20 mM HEPES-KOH pH 7.5, 100 mM KCl). The annealing program consisted of 5 min at 95 °C, followed by a 1 °C/min ramp until 12 °C was reached. The pull-down procedure involved mixing the assembly buffer (40 mM HEPES-KOH pH 7.5, 20 mM MgCl_2_, 50 mM KCl, 2 mM DTT), 300 pmol of DNA substrate, water, and 300 pmol of Cascade-TniQ. The mixture was incubated at room temperature (RT) for 10 min, after which ATP or ATPγS (at 1 mM final concentration) were added to the sample, followed by 2400 pmol of TnsC. The mixture was incubated for 30 min at RT, after which 300 pmol of His -MBP-TnsB^575-613^ or full-length His_6_-MBP-TnsB were added to the sample, reaching a final volume of 53 μL. After 30 min at RT, 147 μL of binding buffer (20 mM HEPES-KOH pH 7.5, 10 mM MgCl_2_, 300 mM KCl, 1 mM DTT, 0.025% tween-20, 1 mM ATP or ATPγS) were added to the sample. 10 μL were withdrawn and mixed with 10 μL of water and 5 μL of SDS dye (225 mM Tris-HCl, pH 6.8, 50% (v/v) glycerol, 5% (w/v) SDS, 0.05% (w/v) bromophenol blue, 250 mM DTT) to obtain a 5% input control sample for SDS-PAGE analysis. The sample was mixed with 50 μL of 50% amylose resin pre-equilibrated in binding buffer and incubated for 1 h at 4 °C on a rotating wheel. Following a short centrifugation (2 min, 4 °C, 500*g*), the supernatant was discarded and the protein-bound resin was washed three times with 500 μL of binding buffer. Ultimately, 25 μL of binding buffer supplemented with 10 mM maltose were added to the sample. After centrifugation (2 minutes at 4 °C, 14000*g*), 20 μL of supernatant were mixed with 5 μL of SDS dye in order to obtain an elution sample for SDS-PAGE analysis. 10 μL of 5% input control and 10 μL of elution sample were loaded on a 10% SDS-PAGE gel (Bio-Rad) for each tested condition. For the control samples Cascade-TniQ-TnsC-minus or MBP-TnsB/AB-minus, the respective SEC buffers were used instead of a protein solution.

### *E. coli* transposition assay

A *Pse*CAST pDonor plasmid was first transformed into *E. coli* BL21 (DE3) cells (NEB). Single colonies were inoculated into liquid culture and the resulting strains were made chemically competent using standard methods. PseCAST pEffector plasmids encoding a crRNA targeting the lacZ locus (crR-NA-4^1^) and all CAST subunits were transformed into the resulting strain and plated on double antibiotic LB-agar plates containing 100 µg/mL carbenicillin and 100 µg/mL spectinomycin. After approximately 20 hours of growth, surviving colonies were scraped from the plates, resuspended in fresh LB medium, and re-plated on double antibiotic LB-agar plates supplemented with 0.1 mM IPTG to induce *Pse*CAST protein expression. Cells were incubated for an additional 18 – 20 hours, scraped, and resuspended in LB medium. Cells were then lysed and integration efficiencies were measured via qPCR as previously described^1^. Wild-type plasmid sequences, crRNA-4, and oligos used for qPCR are listed in **Supplementary Tables 1 and 2**.

### Cascade-TniQ-TnsC complex: cryo-EM sample preparation and data collection

Target and donor substrates were prepared by annealing oligos oMS572, oMS573, and oMS576 (target; **Supplementary Table 3**) and oMS569, oMS570, and oMS571 (donor; **Supplementary Table 3**) at a 1:1:1 molar ratio to 100 μM in annealing buffer (20 mM HEPES-KOH pH 7.5, 100 mM KCl). The annealing program consisted of 5 min at 95 °C followed by a 1 °C/min ramp until 12 °C was reached. For the target pot assembly, 10× buffer (200 mM HEPES-KOH pH 7.5, 1500 mM KCl, 100 mM MgCl_2_, 10 mM DTT), water, and TniQ-Cascade were gently mixed on ice. The target DNA was added and the sample was incubated for 10 min at RT. TnsC and ATP ware added to the sample, which was then incubated for 15 min at RT. The final target pot volume was 25 μL, containing 4.5 μM of Cascade-TniQ, 4.5 μM of target DNA, 40.5 μM of TnsC, 1 mM ATP, 20 mM HEPES-KOH pH 7.5, 150 mM KCl, 10 mM MgCl_2_, and 1 mM DTT. The donor pot was prepared by mixing 10× buffer, water, ATP, TnsAB, and donor DNA to a final volume of 25 μL (containing 18 μM TnsAB, 4.5 μM of donor DNA, 20 mM HEPES-KOH pH 7.5, 150 mM KCl, 10 mM MgCl_2_, 1 mM ATP, and 1 mM DTT) and subsequently incubating the mixture for 10 min at RT. However, a white insoluble precipitation was observed in the donor pot at this stage, and again after mixing with the target pot and incubating at RT for 20 min. This was likely due to TnsAB instability under the conditions used; in fact, the donor pot components were not detected in the later cryo-EM analysis. After mixing the target and donor pots, the sample was centrifuged (14 000*g*, at 4 °C for 10 min) and the pellet discarded. 2.5 μL of supernatant were applied to a freshly glow-discharged (30 s) Quantifoil R 1.2/1.3 Au 300 mesh grid, blotted for 2 s (100% humidity, 4 °C) and plunge-frozen in liquid ethane using a Vitrobot Mark IV plunger (FEI), followed by storage in liquid nitrogen. Cryo-EM data were collected on an FEI Titan Krios G3i (University of Zurich, Switzerland) operating at 300 kV and equipped with a Gatan K3 direct electron detector in super-resolution counting mode. In total, 9403 movies were acquired at ×130,000 magnification, yielding a super-resolution pixel size of 0.325 Å. Each movie consisted of 47 subframes with a cumulative exposure of 59.672 e^-^/Å^2^. EPU Automated Data Acquisition Software for Single Particle Analysis (Thermo Fisher Scientific) was used for data acquisition, with three shots per hole and a defocus range of −1.0 to −2.4 μm in 0.2-μm increments.

### Cascade-TniQ-TnsC complex: cryo-EM data processing and structure modelling

9403 exposures were imported to cryoSPARC^51^ live (v4.3) for real-time processing. After patch motion and patch CTF correction, 745080 particles were identified using blob picker (200-300 Å diameter), extracted (extraction box size: 600 pixels; Fourier-cropped to box size: 200 pixels), and classified into 50 2D classes. 416193 particles were selected and 3 volumes were generated using ab-initio reconstruction. One class yielded a recognizable Cascade-TniQ-TnsC volume (108814 particles) and was used for 3D variability analysis (3DVA) (filter resolution: 5 Å, mode: simple), while the other two volumes did not contain density for TnsC. Two distinct volumes resulting from one 3DVA mode were used as input for a heterogeneous refinement with the aim of separating the particles corresponding to Cascade-TniQ-TnsC from the ones of Cascade-TniQ only. The two volumes, after a round of non-uniform refinement, were used as inputs of a second heterogeneous refinement, together with one of the volumes from the ab-initio multi-class job. This refinement resulted in a volume of Cascade (195798 particles), Cascade-TniQ (143494 particles), and Cascade-TniQ-TnsC (76901 particles). The latter was used for non-uniform refinement after particle re-extraction, yielding a 3.07 Å (GSFSC resolution) reconstruction (76400 particles). After particle subtraction, a local refinement of the TnsC heptamer generated a 3.44 Å (GSFSC resolution) map. A composite map was obtained combining the consensus Cascade-TniQ-TnsC and Tn-sC-focused volumes via the ‘combine_focused_maps’ function in Phenix^52^. Particles corresponding to Cascade only were used as an input for a 3D variability analysis and a heterogeneous refinement in the same way as described above. One of the resulting volumes, corresponding to 132574 particles, yielded a 2.86 Å (GSFSC resolution) Cascade reconstruction after particle re-extraction and non-uniform refinement. The Cascade-TniQ volume was instead fed twice into a heterogeneous refinement. One of the volumes, corresponding to 50074 particles, yielded a 3.19 Å (GSFSC resolution) reconstruction after particle re-extraction and non-uniform refinement. All maps and models have been deposited in the EMDB and PDB databases (see **Data availability** and **Extended Data Table 1**). A detailed processing workflow is shown in the **Extended Data Fig. 1**.

Monomeric initial models of Cas6, Cas7, Cas8, TniQ and TnsC were generated using Alpha-Fold3^53^. The AlphaFold3 models were manually docked as rigid bodies in the cryo-EM density map using UCSF ChimeraX^54^. The model was real-space refined in Phenix against the cryo-EM map and then manually in Coot^55^ using secondary structure, side chain rotamer, Ramachandran, and LibG^56^ nucleic acid restraints. The nucleic acid, comprising the crRNA (positions −7 to 52), target strand (positions −60 to 4) and non-target strand (positions −4 to 4, 21 to 23 and 34 to 60), was manually built in Coot using PDB 7U5D as a starting point. The cryo-EM density was clear enough to distinguish the DNA register. Refinement statistics are reported in **Extended Data Table 1**. Structure and map figures were prepared using UCSF ChimeraX.

### Cascade-TniQ-TnsC-TnsB hook complex: cryo-EM sample preparation and data collection

The cryo-EM sample of the Cascade-TniQ-TnsC-hook complex was prepared using a pull-down approach. A double-stranded target and a double-stranded right-end donor substrate were prepared by annealing oligos oMS628-629 (dsTarget; **Supplementary Table 3**) and oligos oMS576-630 (dsRE; **Supplementary Table 3**) in a 1:1 ratio to a final concentration of 100 μM in annealing buffer (20 mM HEPES-KOH pH 7.5, 100 mM KCl). The annealing program consisted of 5 min at 95 °C followed by a 1 °C/min ramp until 12 °C was reached. The target pot was prepared by adding 1200 pmol of dsTarget to 100 μL of assembly buffer (40 mM HEPES-KOH pH 7.5, 20 mM MgCl_2_, 2 mM DTT), followed by addition of KCl solution to adjust the final KCl concentration to 230 mM, and 1200 pmol of Cascade-TniQ. The mixture was incubated at RT for 10 min, after which ATP and 8400 pmol of TnsC were added. The sample was then incubated at RT for 30 min. The final target pot volume was 200 μL and its buffer composition was 1 mM ATP, 20 mM HEPES-KOH pH 7.5, 10 mM MgCl_2_, 1 mM DTT and 230 mM KCl. A donor pot was prepared by mixing 1200 pmol of His_6_-MBP TnsAB and 1200 pmol of dsRE substrate. The donor pot was brought to a 20 μL final volume using TnsAB SEC buffer (20 mM HEPES-KOH pH 7.5, 500 mM KCl, 1 mM DTT), mixed to the target pot and incubated for 30 min at RT. After adding 30 μL of binding buffer (20 mM HEPES-KOH pH 7.5, 10 mM MgCl_2_, 300 mM KCl, 1 mM DTT, 0.025% tween-20, 1 mM ATP), the sample was mixed with 40 μL of 50% amylose resin pre-equilibrated in binding buffer and incubated for 1 h at 4 °C on a rotating wheel. After centrifugation (2 min, 4 °C, 500*g*), the supernatant was discarded and the protein-bound resin was washed three times with 500 μL of binding buffer. 15 μL of binding buffer supplemented with 10 mM maltose were added and the sample was centrifuged for 5 min at 4 °C, 14000*g*. The supernatant was isolated by careful pipetting and centrifuged two additional times. A second elution was performed using 20 μL of maltose-supplemented binding buffer, followed by three centrifugation steps as described above. The two elutions were combined in one sample and used to freeze cryo-EM grids. 2.5 μL of sample were applied to a freshly glow-discharged (60 s) Quantifoil R 1.2/1.3 Au 300 mesh grid, blotted for 2.5 s at 100% humidity, 4 °C, plunge-frozen in liquid ethane/propane cryogen mixture using a Vitrobot Mark IV plunger (FEI), and stored in liquid nitrogen. Cryo-EM data were collected on an FEI Titan Krios G3i operating at 300 kV and equipped with a Gatan K3 direct electron detector in super-resolution counting mode. In total, 11374 movies were acquired at ×130,000 magnification, yielding a super-resolution pixel size of 0.325 Å. Each movie consisted of 37 subframes with a cumulative exposure of 60.947 e^-^/Å^2^. EPU Automated Data Acquisition Software for Single Particle Analysis (Thermo Fisher Scientific) was used for data acquisition, with three shots per hole and a defocus range of −1.0 to −2.4 μm in 0.2-μm increments.

### Cascade-TniQ-TnsC-TnsB hook complex: cryo-EM data processing and structure modelling

11374 exposures were imported to cryoSPARC live (v4.7) for real-time processing. After patch motion and patch CTF correction, 329059 particles were identified using blob picker (200-300 Å diameter), extracted (extraction box size: 600 pixels; Fourier-cropped to box size: 200 pixels), and classified into 50 2D classes. The selected 114608 particles were used as a training set for Topaz picking (350 Å expected particle diameter, 50 particles per micrograph), yielding 463242 particles, which were extracted (extraction box size: 500 pixels; Fourier-cropped to box size: 250 pixels) and classified in 150 2D classes. Classes corresponding to 189451 particles were selected and fed into a 2-classes ab-initio reconstruction, resulting in a junk class and a recognizable Cascade-TniQ-TnsC volume (142343 particles), which was used as input for non-uniform refinement after particle re-extraction. A 2.84 Å (GSFSC resolution) reconstruction was obtained from a final number of 138834 particles, which was further used for a TnsC-focused local refinement, yielding a 2.96 Å (GSFSC resolution) volume of the heptamer decorated with additional density for each of its seven subunits. A composite map was obtained combining the consensus Cascade-TniQ-TnsC and TnsC-focused volumes via the ‘combine_focused_maps’ function in Phenix. All maps and models have been deposited in the EMDB and PDB databases (see **Data availability** and **Extended Data Table 1**). A detailed processing workflow is shown in the **Extended Data Fig. 5**.

The previously obtained Cascade-TniQ-TnsC model was rigid-body fitted in the cryo-EM density map using UCSF ChimeraX and manually adjusted in Coot using the above-mentioned restraints. The target and non-target strand DNA were extended by three bases at the 5’ TS-end and by two bases at the 3’ TS-end. A multimer of the TnsC heptamer bound to seven TnsB^575-613^ copies was generated using AlphaFold3. The model positioned the last nine TnsB C-terminal residues at the additional densities of each TnsC subunit. Although the map resolution worsens radially from the centre of the protein ring (**Extended Data Fig. 5c**), the recognizable density of the bulky residues W605 and F612 aided in positioning the peptide in the density. Refinement statistics are reported In **Extended Data Table 1**. Structure and map figures were prepared using UCSF ChimeraX.

### PseCAST transpososome holocomplex: cryo-EM sample preparation and data collection

The cryo-EM sample of the Cascade-TniQ-TnsC-TnsAB holocomplex was prepared using a pulldown approach. The strand-transfer substrate was prepared by annealing oligos oMS733, oMS734 and oSO146 (**Supplementary Table 3**) in a 1:1:1 ratio and to a final concentration of 100 μM each. The annealing program consisted of 5 min at 95 °C, followed by a 1 °C/min ramp until 12 °C was reached. 1000 pmol of substrate, 22.5 μL of assembly buffer A (40 mM HEPES-KOH pH 7.5, 20 mM MgCl_2_, 200 mM KCl, 2 mM DTT), and water were mixed with 2000 pmol of His_6_-MBP-TnsAB in a final volume of 45 μL. The sample was kept on ice for 10 min. In the meantime, 35.8 μL (1000 pmol) of Cascade-TniQ were mixed with 55 μL of assembly buffer B (40 mM HEPES-KOH pH 7.5, 20 mM MgCl_2_, 100 mM KCl, 2 mM DTT) and 1.6 μL of water in order to lower its KCl concentration to 260 mM. This mixture was added to the initial pot and incubated for 10 min at RT. ATP (1 mM final concentration) was added, followed by 7000 pmol of TnsC, thereby reaching a final volume of 155 μL. The mixture was then incubated at RT for 30 min. After the addition of 45 μL of binding buffer (20 mM HEPES-KOH pH 7.5, 10 mM MgCl_2_, 300 mM KCl, 1 mM DTT, 0.025% tween-20, 1 mM ATP), the sample was mixed with 50 μL of 50% amylose resin pre-equilibrated in binding buffer. The mixture was incubated for 1.5 h at 4 °C on a rotating wheel and, following centrifugation (2 min, 4 °C, 500*g*), the supernatant was discarded and the protein-bound resin was washed three times with 500 μL of binding buffer. After adding 25 μL of binding buffer supplemented with 10 mM maltose, the sample was centrifuged for 10 minutes at 4 °C, 14000*g*. The supernatant was isolated by careful pipetting and centrifuged two additional times. 2.5 μL of sample were applied to a freshly glow-discharged (60 s) Quantifoil R 1.2/1.3 Au 300 mesh grid, blotted for 3 (grid 1) or 2.5 s (grid 2) at 100% humidity and 4 °C, plunge-frozen in a liquid ethane/propane cryogenic mixture using a Vitrobot Mark IV plunger (FEI), and stored in liquid nitrogen. Cryo-EM data were collected on an FEI Titan Krios G3i operating at 300 kV and equipped with a Gatan K3 direct electron detector in super-resolution counting mode. In total, 43220 movies were acquired over two separate data collections (dataset 1, 11964 movies of grid 1; and dataset 2, 31256 movies of grid 2) at ×130,000 magnification, yielding a super-resolution pixel size of 0.325 Å. Each movie consisted of 37 subframes with a cumulative exposure of 60.059 (dataset 1) or 65.580 (dataset 2) e^-^/Å^2^. EPU Automated Data Acquisition Software for Single Particle Analysis (Thermo Fisher Scientific) was used for data acquisition, with three shots per hole and a defocus range of −1.0 to −2.4 μm in 0.2-μm increments.

### PseCAST transpososome holocomplex: cryo-EM data processing and structure modelling

For both data collections, the exposures were imported to cryoSPARC live (v4.7) for real-time processing. Dataset 1 and dataset 2 were pre-processed separately. After patch motion and patch CTF correction, particles were identified using blob picker (270-290 Å diameter, elliptical blob), extracted (extraction box size: 600 pixels; Fourier-cropped to box size: 200 pixels), and classified into 100 2D classes. 93462 particles of dataset 1 and 203735 of dataset 2 were selected for a 2-classes ab-initio reconstruction, that resulted in a Cascade-TniQ-TnsC and a TnsAB tetramer volumes. The Cascade-TniQ-TnsC particles were re-extracted with a bigger box (1024/256 px) and used for non-uniform refinement. Using a bigger box revealed additional density at the edge of the particles, corresponding to a flexible TnsAB tetramer next to the TnsC density. This volume was low-pass filtered and used to create masks for particle subtraction. After reducing the box size (700/350 px), the signal for Cascade-TniQ was subtracted from the selected set of particles and the TnsABC moiety was locally refined after aligning the volume to the centre of mass of the TnsABC mask. 3D variability analysis (filter resolution: 5 Å; mode: simple) yielded one reconstruction with clear density for both TnsC and TnsAB, and another with well-resolved TnsC but only weak density for TnsAB. The volumes were used as input for two rounds of heterogeneous refinement with the initial set of particles. 49752 were assigned to the TnsAB-bound TnsC volume, re-extracted (576 px) and locally refined focusing on TnsAB. In order to adjust the box size for the creation of a composite map, the particles were re-extracted with a larger box-size (700 px) and re-aligned to the centre of mass of the Cascade-TniQ-TnsC mask, yielding a 49606 particle set. An ab-initio job, followed by non-uniform refinement, resulted in a Cascade-TniQ-TnsC-TnsAB consensus volume at 3.10 Å GSFSC resolution. Using the consensus map, new masks were generated for the Cascade-TniQ-TnsC, TnsC and TnsAB moieties as described above. The masks were used for particle subtraction of the TnsAB signal and local refinement of the Cascade-TniQ-TnsC complex. To improve the signal of the TnsB-hooks bound to TnsC, a final particle subtraction of the Cascade-TniQ signal was followed by a local refinement of TnsC. The TnsAB local refinement was repeated using the final set of 49606 particles in order to match the rest of the volumes. This procedure yielded focused cryo-EM maps for the TnsAB tetramer, the hook-bound TnsC heptamer, and the Cascade-TniQ-TnsC-hook complex at a 3.36 Å, 3.30 Å and 3.09 Å GSFSC resolution, respectively. After re-alignment of the TnsAB volume, the global consensus map and the locally refined maps were finally combined into a composite map using the ‘volume maximum’ function of ChimeraX. All maps and models have been deposited in the EMDB and PDB databases (see **Data availability** and **Extended Data Table 1**). A detailed processing workflow is shown in the **Extended Data Fig. 8**.

The previously obtained Cascade-TniQ-TnsC-hook model was rigid-body fitted in the cryo-EM density map using UCSF ChimeraX and manually adjusted in Coot using the above-mentioned restraints. The density corresponding to the tetrameric TnsAB was rigid-body fitted using our previously determined TnsAB PEC structure (PDB 9T7L). Missing regions in the protein and nucleic acid chains were manually built in Coot with the aid of an AlphaFold3 model of TnsAB. It was not possible to determine whether the hooks bound to TnsC could be assigned to any of the TnsB copies observed in the structure, as the region between the hook and the last modelled TnsB residue (TnsB 586-604) is disordered. It should also be noted that the 7:7 TnsC:hook observed stoichiometry likely reflects the excess of bait (His_6_-MBP-TnsAB) used in pull-down preparation, rather than the physiological interaction ratio, which was observed to be 7:4 in other transpososome structures^6,36,41^. Refinement statistics are reported in **Extended Data Table 1**. Structure and map figures were prepared using UCSF ChimeraX.

## DATA AVAILABILITY

Cryo-EM reconstructions have been deposited in the Electron Microscopy Data Bank under accession codes EMD-57765, EMD-57750, EMD-57751, EMD-57757, EMD-57758 (*Pse*Cascade-TniQ-TnsC complex and related maps), EMD-57737, EMD-57738, EMD-57739 (*Pse*Cascade-TniQ-TnsC complex bound to TnsB-hook motifs and related maps) and EMD-57728, EMD-57729, EMD-57730, EMD-57731, EMD-57736 (*Pse*Cascade-TniQ-TnsC-TnsAB holocomplex and related maps). Coordinates for atomic models have been deposited in the Protein Data Bank under accession codes 30GT (*Pse*Cascade-TniQ-TnsC complex), 30GB (*Pse*Cascade-TniQ-TnsC complex bound to TnsB-hook motifs) and 30GA (*Pse*Cascade-TniQ-TnsC-TnsAB holocomplex).

## ACKNOWLEDGEMENTS

We are grateful to the staff at UZH Center for Microscopy and Image Analysis for providing electron microscope access and technical support. We thank L. Ronner for technical assistance, T.M. Smith for laboratory support, L. F. Landweber for qPCR instrument access, and the JP Sulzberger Columbia Genome Center for NGS support. We thank members of the Jinek and Sternberg labs for discussions and critical feedback on the study.

## AUTHOR CONTRIBUTIONS

G.F., M.S., S.O, M.J. and S.H.S. conceived the study and designed experiments. M.J. and S.H.S. supervised the study. S.O. and M.S. prepared cryo-EM samples and collected cryo-EM data of the *Pse-* Cascade-TniQ-TnsC complex. G.F. prepared cryo-EM samples and collected cryo-EM data of the *Pse-* Cascade-TniQ-TnsC complex bound to TnsB-hook motifs and the *Pse*Cascade-TniQ-TnsC-TnsAB holocomplex. G.F. processed all cryo-EM data and built the structural models together with S.O. G.D.L. carried out *in vivo* transposition assays in *E. coli*. G.F., M.J., and S.O. analysed data. G.F., S.O., M.J. and S.H.S. wrote the manuscript with input from G.D.L., M.S..

