## Extended Data for "Structural basis of RNA-guided DNA integration by type I CRISPR-associated transposases"

### EXTENDED DATA TABLES AND FIGURES

#### **Extended Data Table 1** Cryo-EM data collection, refinement and validation statistics

| ID, EMDB and PDB codeS | Cryo-EM structure of the *Pse*Cas-cade-TniQ-TnsC complex<br>(EMD-57765)<br>(PDB-30GT) | Consensus cryo-EM volume of the *Pse*-Cascade-TniQ-TnsC complex<br>(EMD-57751) | TnsC-focused cryo-EM volume of the *Pse*Cas-cade-TniQ-TnsC complex<br>(EMD-57750) | Cryo-EM volume of the *Pse*Cas-cade-TniQ complex<br>(EMD-57757) | Cryo-EM volume of the *Pse*Cascade complex<br>(EMD-57758) |
|---|---|---|---|---|---|
| **Data collection and processing** | | | | | |
| Magnification | 130,000 | 130,000 | 130,000 | 130,000 | 130,000 |
| Voltage (kV) | 300 | 300 | 300 | 300 | 300 |
| Electron exposure ($e^-$ /Å$^2$) | 59.672 | 59.672 | 59.672 | 59.672 | 59.672 |
| Defocus range (μm) | -1.0 to -2.4 (0.2 steps) | -1.0 to -2.4 (0.2 steps) | -1.0 to -2.4 (0.2 steps) | -1.0 to -2.4 (0.2 steps) | -1.0 to -2.4 (0.2 steps) |
| Pixel size (Å) | 0.325 | 0.325 | 0.325 | 0.325 | 0.325 |
| Symmetry imposed | C1 | C1 | C1 | C1 | C1 |
| Initial particle images (no.) | 745,080 | 745,080 | 745,080 | 745,080 | 745,080 |
| Final particle images (no.) | 76,400 | 76,400 | 76,400 | 50,074 | 132,574 |
| Map resolution (Å) | 3.26 (composite) | 3.07 | 3.44 | 3.19 | 2.86 |
| FSC threshold | 0.143 | 0.143 | 0.143 | 0.143 | 0.143 |
| Map resolution range (Å) | n/a | 2.6-6.0 | 3.0-7.0 | 2.5-8.0 | 2.5-6.0 |
| **Refinement** | | | | | |
| Initial model used (PDB code) | AlphaFold3, 7U5D | n/a | n/a | n/a | n/a |
| Model resolution (Å) | 3.0 | n/a | n/a | n/a | n/a |
| FSC threshold | 0.143 | n/a | n/a | n/a | n/a |
| Model resolution range (Å) | 3.0-3.4 | n/a | n/a | n/a | n/a |
| Map sharpening *B* factor (Å$^2$) | n/a | 58.0 | 65.2 | 52.6 | 71.6 |
| Model composition<br>Non-hydrogen atoms<br>Protein residues<br>Nucleotide residues<br>Ligands | <br>51,759<br>5,959<br>165<br>ATP: 7, MG: 7 | n/a | n/a | n/a | n/a |
| *B* factors (Å$^2$) min/max/mean<br>Protein<br>Nucleotide<br>Ligand | <br>53.14/241.07/115.81<br>72.66/309.92/143.87<br>71.01/148.12/104.08 | n/a | n/a | n/a | n/a |
| R.m.s. deviations<br>Bond lengths (Å)<br>Bond angles (°) | <br>0.006(0)<br>0.564(0) | n/a | n/a | n/a | n/a |
| Validation<br>MolProbity score<br>Clashscore<br>Poor rotamers (%) | <br>1.86<br>8.16<br>2.47 | n/a | n/a | n/a | n/a |
| Ramachandran plot<br>Favored (%)<br>Allowed (%)<br>Disallowed (%) | <br>0.00<br>2.60<br>97.40 | n/a | n/a | n/a | n/a |

| ID, EMDB and PDB codes | Cryo-EM structure of the *Pse*Cascade-TniQ-TnsC complex bound to *Pse*TnsB-hook motifs<br>(EMD-57739)<br>(PDB-30GB) | Consensus cryo-EM volume of the *Pse*Cascade-TniQ-TnsC complex bound to TnsB-hook motifs<br>(EMD-57738) | TnsC-focused cryo-EM volume of the *Pse*Cascade-TniQ-TnsC complex bound to TnsB-hook motifs<br>(EMD-57737) |
|---|---|---|---|
| **Data collection and processing** | | | |
| Magnification | 130,000 | 130,000 | 130,000 |
| Voltage (kV) | 300 | 300 | 300 |
| Electron exposure ($e^-$/Å$^2$) | 60.947 | 60.947 | 60.947 |
| Defocus range (μm) | -1.0 to -2.4 (0.2 steps) | -1.0 to -2.4 (0.2 steps) | -1.0 to -2.4 (0.2 steps) |
| Pixel size (Å) | 0.325 | 0.325 | 0.325 |
| Symmetry imposed | C1 | C1 | C1 |
| Initial particle images (no.) | 329,059 | 329,059 | 329,059 |
| Final particle images (no.) | 138,834 | 138,834 | 138,834 |
| Map resolution (Å) | 2.9 (composite) | 2.84 | 2.96 |
| FSC threshold | 0.143 | 0.143 | 0.143 |
| Map resolution range (Å) | n/a | 2.5-5.0 | 2.5-5.0 |
| **Refinement** | | | |
| Initial model used (PDB code) | 30GT, AlphaFold3 | n/a | n/a |
| Model resolution (Å) | 2.8 | n/a | n/a |
| FSC threshold | 0.143 | n/a | n/a |
| Model resolution range (Å) | 2.8-3.2 | n/a | n/a |
| Map sharpening *B* factor (Å$^2$) | n/a | 72.7 | 83.7 |
| Model composition | | n/a | n/a |
| Non-hydrogen atoms | 52,521 | | |
| Protein residues | 6,022 | | |
| Nucleotide residues | 175 | | |
| Ligands | ATP: 7, MG: 7 | | |
| *B* factors (Å$^2$) min/max/mean | | n/a | n/a |
| Protein | 56.32/220.70/112.76 | | |
| Nucleotide | 76.75/313.01/147.48 | | |
| Ligand | 68.42/103.32/83.71 | | |
| R.m.s. deviations | | n/a | n/a |
| Bond lengths (Å) | 0.003(0) | | |
| Bond angles (°) | 0.570(0) | | |
| Validation | | n/a | n/a |
| MolProbity score | 1.70 | | |
| Clashscore | 7.32 | | |
| Poor rotamers (%) | 1.83 | | |
| Ramachandran plot | | n/a | n/a |
| Favored (%) | 0.00 | | |
| Allowed (%) | 2.47 | | |
| Disallowed (%) | 97.53 | | |

| ID, EMDB and PDB codes | Cryo-EM structure of the PseCascade-TniQ-TnsC-TnsAB holocomplex (EMD-57736) (PDB-30GA) | Consensus cryo-EM volume of the PseCascade-TniQ-TnsC-TnsAB holocomplex (EMD-57731) | Cascade-TniQ-TnsC-focused cryo-EM volume of the PseCascade-TniQ-TnsC-TnsAB holocomplex (EMD-57730) | TnsC-focused cryo-EM volume of the PseCascade-TniQ-TnsC-TnsAB holocomplex (EMD-57729) | TnsAB-focused cryo-EM volume of the PseCascade-TniQ-TnsC-TnsAB holocomplex (EMD-57728) |
|---|---|---|---|---|---|
| **Data collection and processing** | | | | | |
| Magnification | 130,000 | 130,000 | 130,000 | 130,000 | 130,000 |
| Voltage (kV) | 300 | 300 | 300 | 300 | 300 |
| Electron exposure ($e^-$ /Å$^2$) | 60.059 (grid1) / 65.580 (grid2) | 60.059 (grid1) / 65.580 (grid2) | 60.059 (grid1) / 65.580 (grid2) | 60.059 (grid1) / 65.580 (grid2) | 60.059 (grid1) / 65.580 (grid2) |
| Defocus range (μm) | -1.0 to -2.4 (0.2 steps) | -1.0 to -2.4 (0.2 steps) | -1.0 to -2.4 (0.2 steps) | -1.0 to -2.4 (0.2 steps) | -1.0 to -2.4 (0.2 steps) |
| Pixel size (Å) | 0.325 | 0.325 | 0.325 | 0.325 | 0.325 |
| Symmetry imposed | C1 | C1 | C1 | C1 | C1 |
| Initial particle images (no.) | 1,919,871 | 1,919,871 | 1,919,871 | 1,919,871 | 1,919,871 |
| Final particle images (no.) | 49,606 | 49,606 | 49,606 | 49,606 | 49,606 |
| Map resolution (Å) | 3.21 (composite) | 3.10 | 3.09 | 3.30 | 3.36 |
| FSC threshold | 0.143 | 0.143 | 0.143 | 0.143 | 0.143 |
| Map resolution range (Å) | n/a | 2.5-8.5 | 2.5-6.5 | 2.5-6.5 | 2.5-6.5 |
| **Refinement** | | | | | |
| Initial model used (PDB code) | 30GB, 9T7L, AlphaFold3 | n/a | n/a | n/a | n/a |
| Model resolution (Å) | 2.8 | n/a | n/a | n/a | n/a |
| FSC threshold | 0.143 | n/a | n/a | n/a | n/a |
| Model resolution range (Å) | 2.1-3.5 | n/a | n/a | n/a | n/a |
| Map sharpening *B* factor (Å$^2$) | n/a | 43.1 | 35.9 | 50.8 | 52.3 |
| Model composition | | n/a | n/a | n/a | n/a |
| Non-hydrogen atoms | 74407 | | | | |
| Protein residues | 8274 | | | | |
| Nucleotide residues | 348 | | | | |
| Ligands | ATP: 7, MG: 13 | | | | |
| *B* factors (Å$^2$) min/max/mean | | n/a | n/a | n/a | n/a |
| Protein | 30.00/270.24/109.72 | | | | |
| Nucleotide | 20.00/323.94/148.85 | | | | |
| Ligand | 68.21/155.99/89.75 | | | | |
| R.m.s. deviations | | n/a | n/a | n/a | n/a |
| Bond lengths (Å) | 0.004(0) | | | | |
| Bond angles (°) | 0.617(0) | | | | |
| Validation | | n/a | n/a | n/a | n/a |
| MolProbity score | 1.87 | | | | |
| Clashscore | 8.51 | | | | |
| Poor rotamers (%) | 2.19 | | | | |
| Ramachandran plot | | n/a | n/a | n/a | n/a |
| Favored (%) | 0.00 | | | | |
| Allowed (%) | 2.83 | | | | |
| Disallowed (%) | 97.17 | | | | |

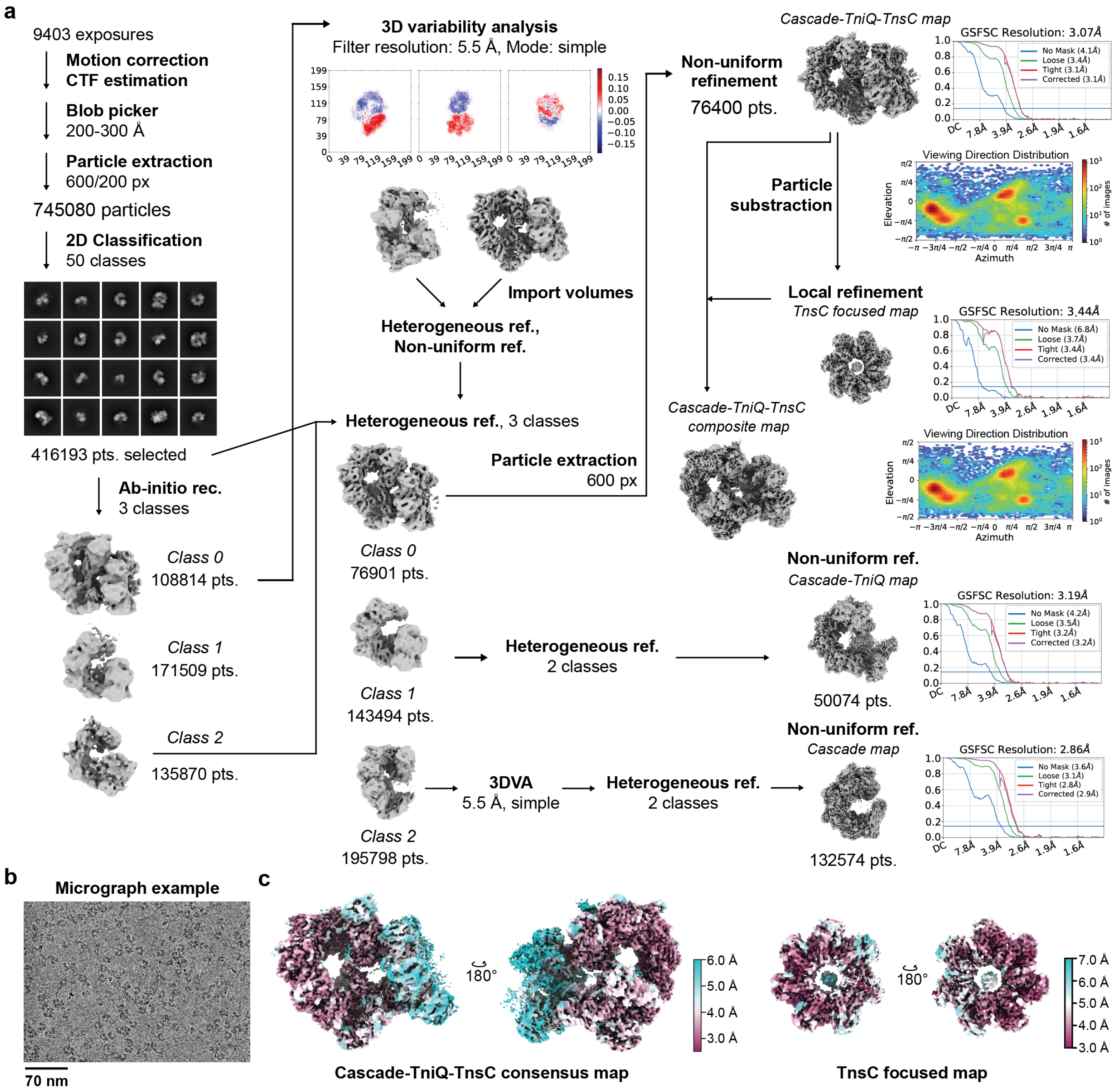


**Extended Data Fig. 1. Cryo-EM data processing of the *Pse*Cascade-TniQ-TnsC targeting complex. a**, Cryo-EM image processing workflow for the *Pse*Cascade-TniQ-TnsC complex. Resolution is determined by Fourier Shell Correlation (FSC) calculated from two independently refined half-maps. The gold-standard cutoff (FSC = 0.143) is marked with a blue line. Final maps were obtained for *Pse*Cascade-TniQ-TnsC (top, 3.07 Å), *Pse*Cascade-TniQ (middle, 3.19 Å), and *Pse*Cascade (bottom, 2.86 Å). **b**, Representative micrograph. **c**, Final consensus and locally refined electron density maps for *Pse*Cascade-TniQ-TnsC and TnsC only, coloured according to local resolution.

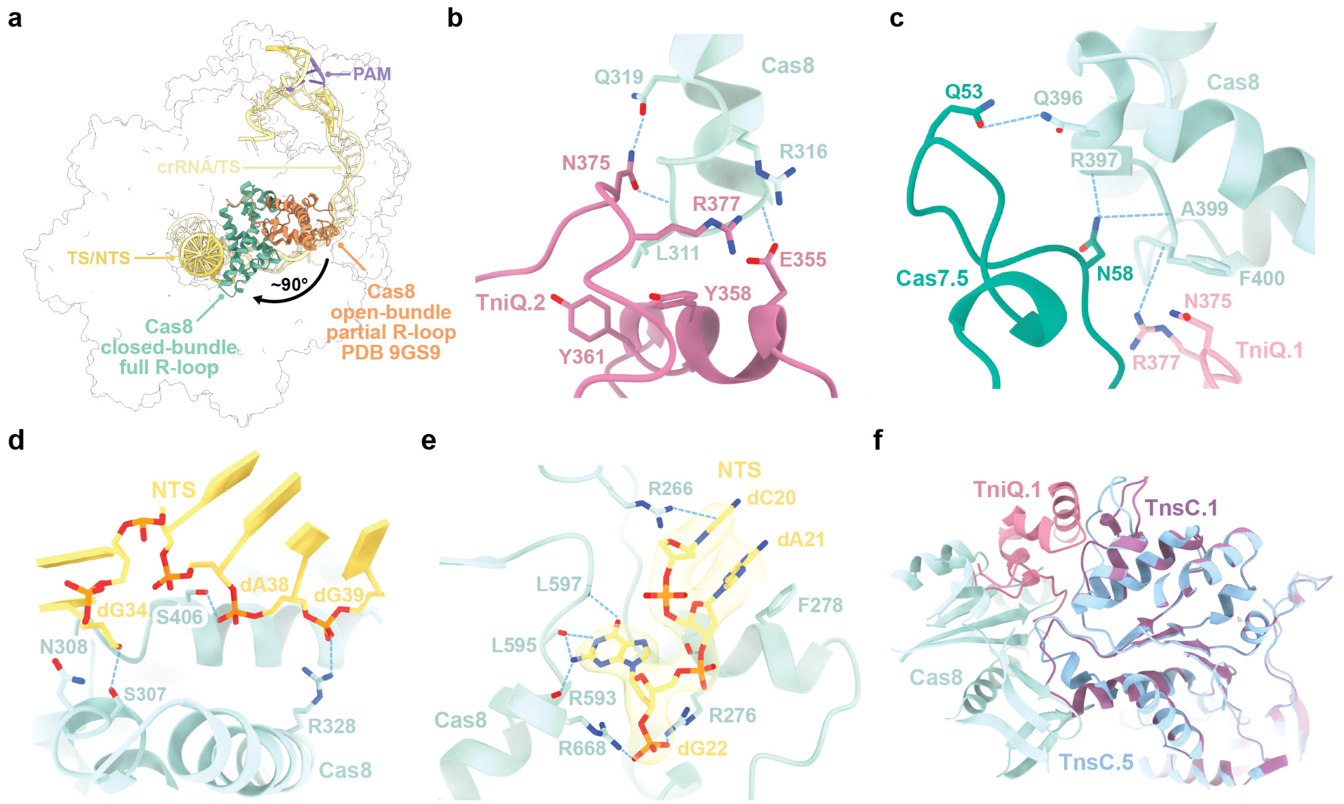


**Extended Data Fig. 2. Domain rearrangements and interaction interfaces of Cas8. a**, Rotation of the Cas8 bundle in the *Pse*Cascade complex with respect to its partial R-loop state (PDB 9GS9)[9]. **b-d**, Structural details of the interactions of the Cas8 helical bundle with TniQ.2 (**b**), TniQ.1 and Cas7.5 (**c**), and the PAM-distal non-target strand (NTS) (**d**). **e**, Binding of the displaced NTS by the Cas8 bundle. Bases are shown in stick representation, with corresponding cryo-EM density (transparent surface in yellow) superimposed . **f**, Superposition of TnsC.1-TniQ and TnsC.5-Cas8 interfaces.

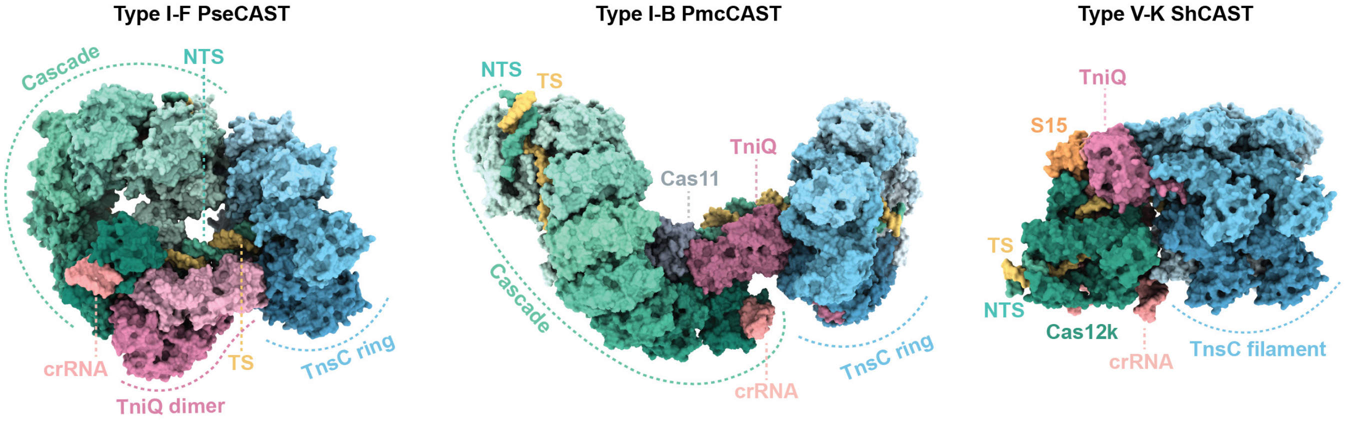


**Extended Data Fig. 3. Structural comparisons of targeting complexes from representative CAST systems.** Side-by-side views of type I-F *Pse*Cascade-TniQ-TnsC (this study), type I-B *Pmc*Cascade-TniQ-TnsC (PDB 8FF4)[41], and type V-K *Sh*Cas12k-S15-TniQ-TnsC (PDB 8EA3)[6]; components beyond the TnsC filament were removed for clarity). Analogous components are highlighted in the same colours across systems, and structures are aligned using the matchmaker function in ChimeraX.

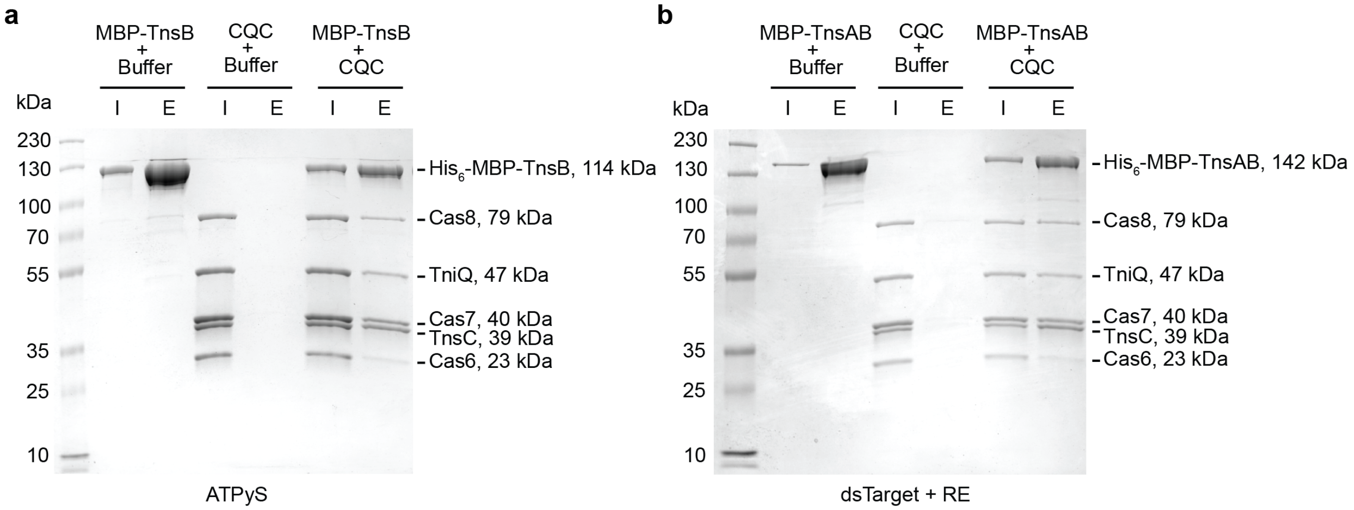


**Extended Data Fig. 4. Co-precipitation of *Pse*Cascade-TniQ-TnsC by TnsB or TnsAB. a**, Co-precipitation of Cascade-TniQ-TnsC by full-length His$_6$-MBP-TnsB in the presence of ATPγS and double-stranded target DNA (dsTarget). **b**, Reconstitution of the *Pse*Cascade-TniQ-TnsC-TnsB-hook complex for cryo-EM analysis. The complex was assembled using co-precipitation of dsTarget DNA-bound Cascade-TnsC-TniQ by amylose-immobilized full-length TnsAB bound to double stranded transposon right end (RE) DNA in the presence of ATP. CQC, Cascade-TniQ-TnsC; I, 5% input control; E, elution.

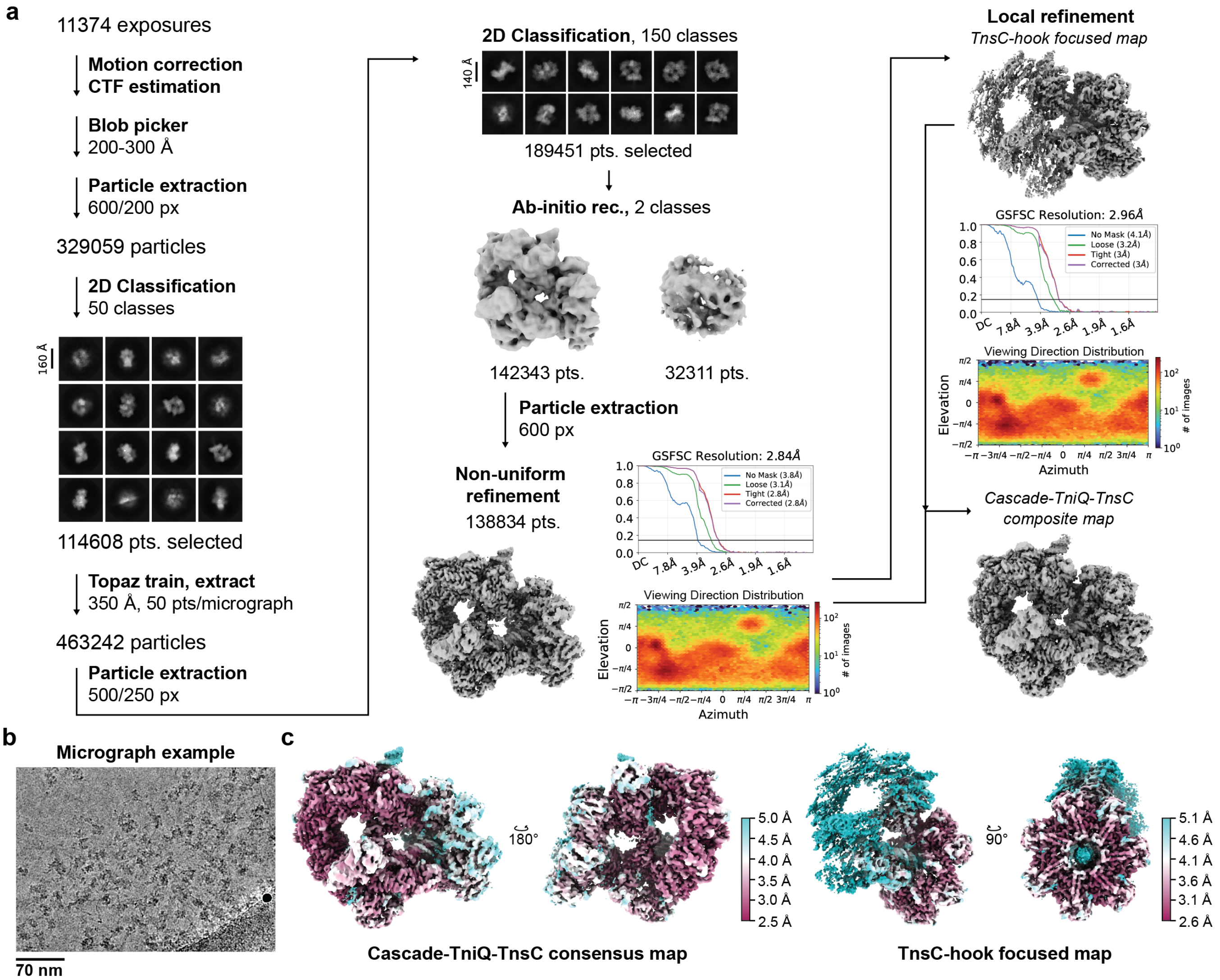


**Extended Data Fig. 5. Cryo-EM data processing of the *Pse*Cascade-TniQ-TnsC-TnsB-transposase recruitment complex. a**, Cryo-EM image processing workflow for the *Pse*Cascade-TniQ-TnsC-TnsB-hook complex. Resolution is determined by Fourier Shell Correlation (FSC) calculated from two independently refined half-maps. The gold-standard cutoff (FSC = 0.143) is marked with a black line. **b**, Representative micrograph. **c**, Final consensus and locally refined electron density maps for *Pse*Cascade-TniQ-TnsC-TnsB-hook and TnsC-hook only, coloured according to local resolution.

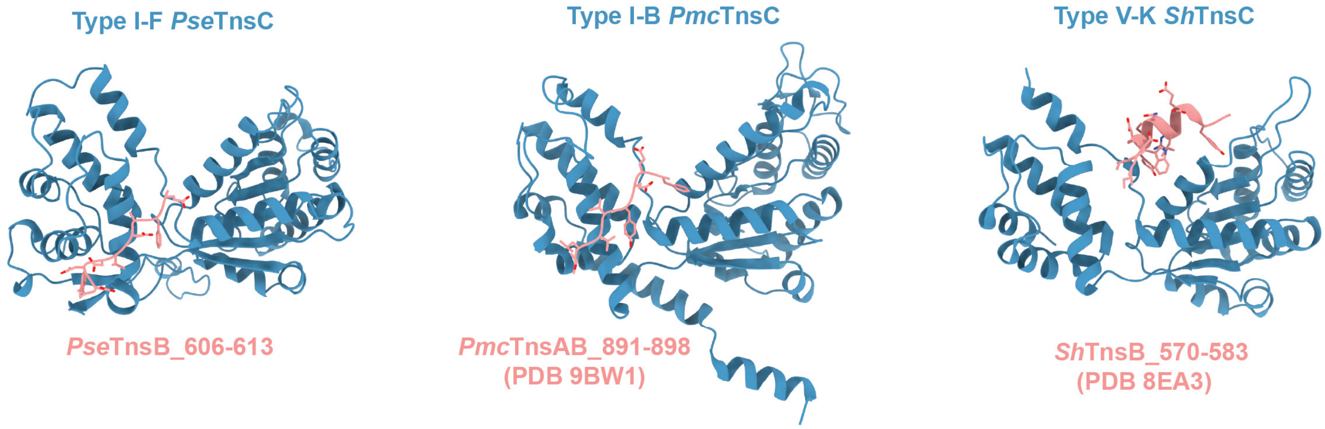


**Data Fig. 6. Comparison of TnsB hook-bound TnsC structures.** Side-by-side views of the TnsC-TnsB hook interfaces in type I-F *Pse*CAST (this study), type I-B *Pmc*CAST (PDB 9BW1), and type V-K *Sh*CAST (PDB 8EA3), shown in the same orientation.

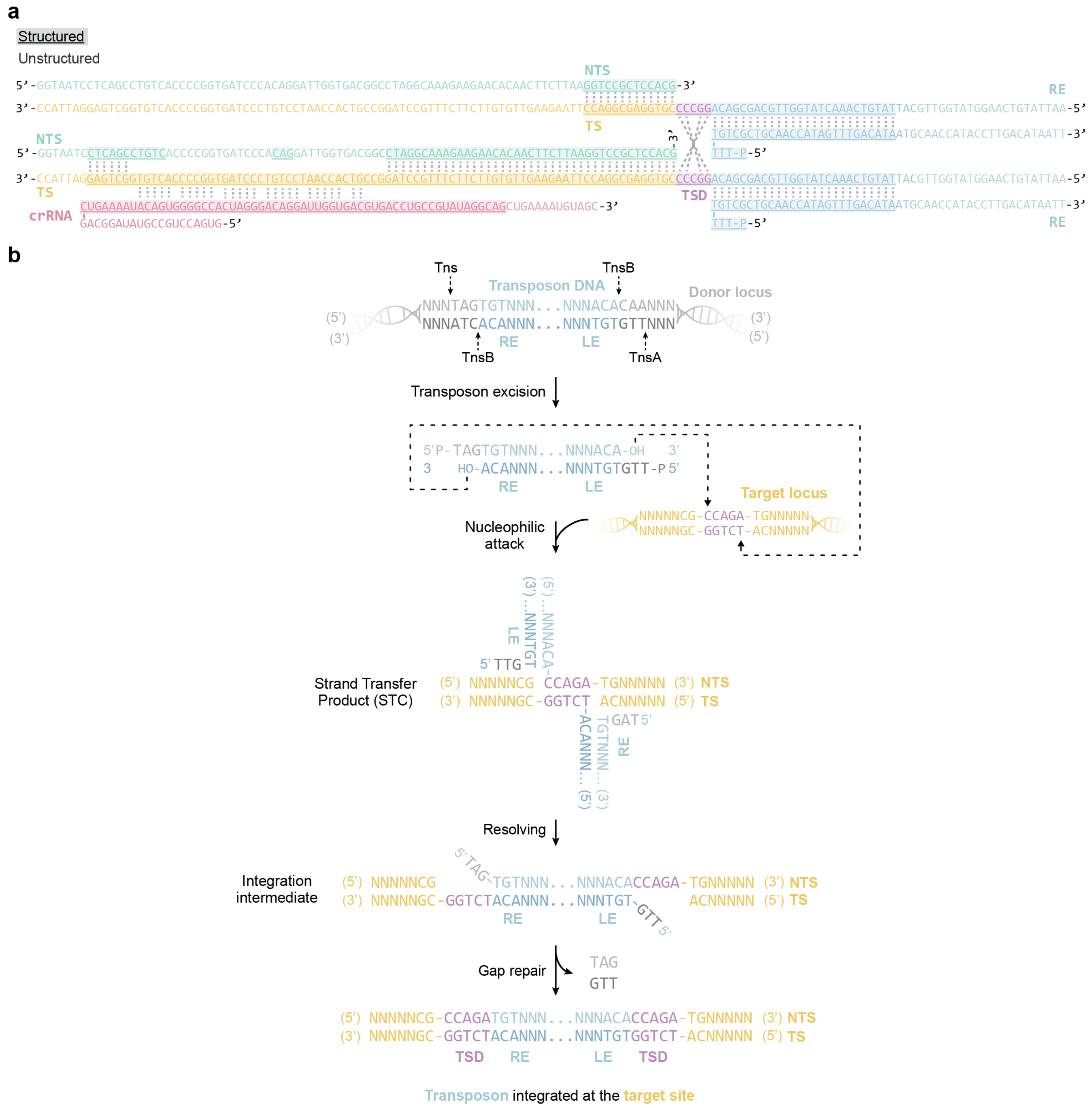


**Extended Data Fig. 7. Transposition DNA constructs. a**, Full-length sequence and base pairing of the oligonucleotides used for *Pse*CAST holocomplex reconstitution, coloured as in **Fig. 3a**. The structured portion modelled in the structure is highlighted. **b**, Sequence-level schematic diagram of the donor and target DNAs and the cut-and-paste transposition cycle of *Pse*CAST. The transposases TnsA and TnsB recognize and process the left (LE) and right (RE) ends, excising the transposon DNA, which is subsequently integrated at the target site through a strand-transfer intermediate by TnsB. TS: target strand; NTS: non-target strand; TSD: target site duplication.

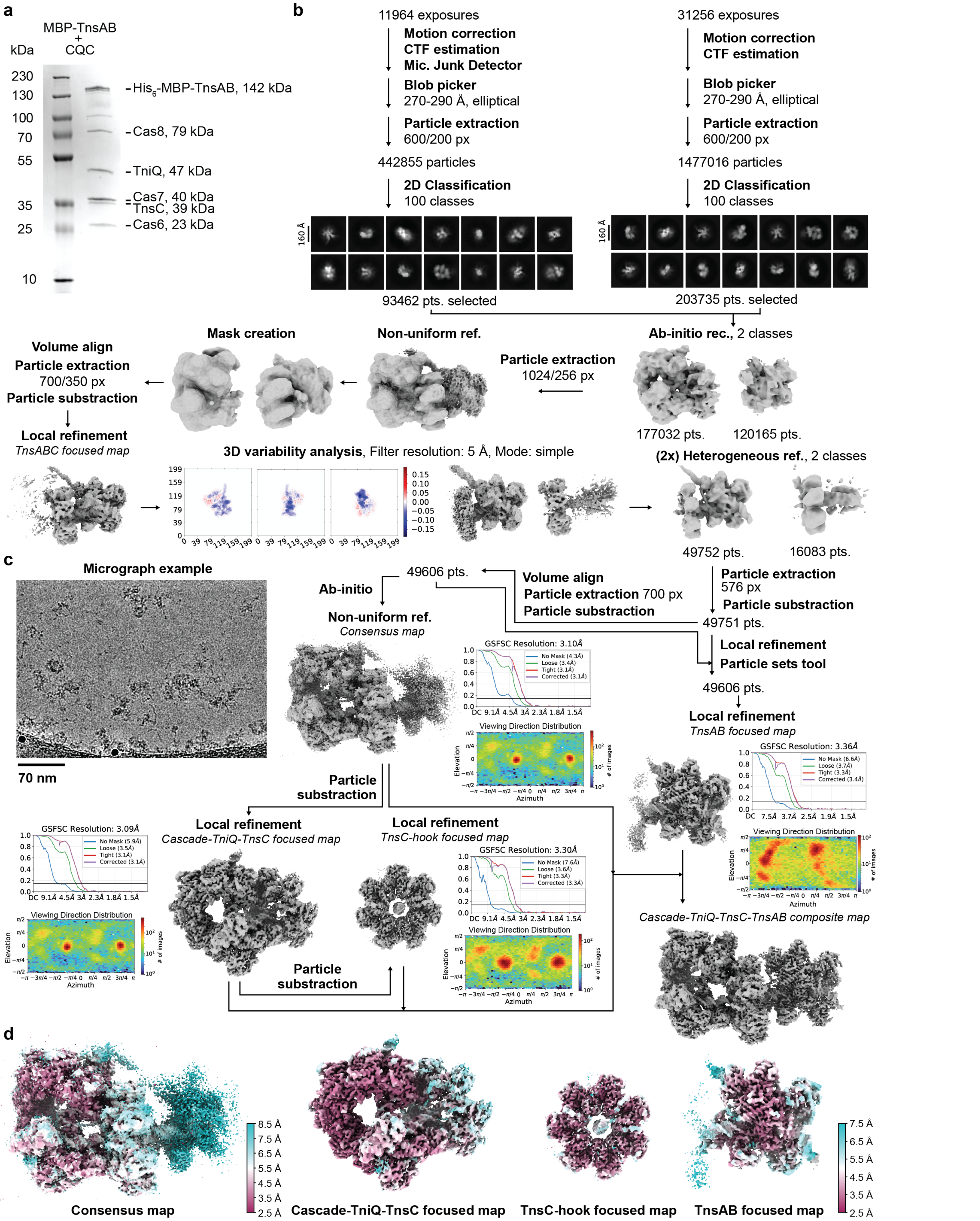


**Extended Data Fig. 8. Cryo-EM sample preparation and data processing of the *Pse*Cascade-TniQ-TnsC-TnsAB holocomplex. a**, SDS-PAGE of the eluate from the co-precipitation of *Pse*Cascade-TnsC-TniQ by amylose-immobilized full-length $His_6$-MBP-tagged *Pse*TnsAB bound to a strand-transfer DNA construct containing a target site, a pseudo-palindromic TSD and a double-stranded RE. The sample was used for cryo-EM analysis to obtain the *Pse*Cascade-TniQ-TnsC-TnsAB holocomplex structure. **b**, Cryo-EM image processing workflow for the *Pse*Cascade-TniQ-TnsC-TnsAB holocomplex. Resolution is determined by Fourier Shell Correlation (FSC) calculated from two independently refined half-maps. The gold-standard cutoff (FSC = 0.143) is marked with a black line. **c**, Representative micrograph. **d**, Global consensus electron density map of the *Pse*Cascade-TniQ-TnsC-TnsAB complex and locally refined volumes of the *Pse*Cascade-TniQ-TnsC, TnsC-hook, and TnsAB subcomplexes, coloured according to local resolution.
